# SARS-CoV-2 Variant Delta Potently Suppresses Innate Immune Response and Evades Interferon-Activated Antiviral Responses

**DOI:** 10.1101/2022.02.22.481430

**Authors:** Dixit Tandel, Vishal Sah, Nitesh Kumar Singh, Poojitha Sai Potharaju, Divya Gupta, Sauhard Shrivastava, Divya Tej Sowpati, Krishnan H Harshan

## Abstract

Delta variant of SARS-CoV-2 has caused more severe infections than its previous variants. We studied the host innate immune response to Delta, Alpha and two earlier variants to map the evolution of the recent ones. Our biochemical and transcriptomic studies reveal that Alpha and Delta have progressively evolved over the ancestral variants by silencing innate immune response, thereby limiting cytokine and chemokine production. Though Alpha silenced RLR pathway just as Delta, it failed to persistently silence the innate immune response unlike Delta. Both Alpha and Delta have evolved to resist IFN treatment while they are still susceptible to RLR activation, further highlighting the importance of RLR-mediated, IFN-independent mechanisms in restricting SARS-CoV-2. Our studies reveal that SARS-CoV-2 Delta has integrated multiple mechanisms to silence host innate immune response and evade IFN response. Delta’s silent replication and sustained suppression of host innate immune response, possibly resulting in delayed or reduced intervention by the adaptive immune response, could potentially contribute to the severe symptoms and poor recovery index associated with it.

## INTRODUCTION

SARS-CoV-2, the causative virus behind the current COVID-19 pandemic has been evolving since its first detection in humans in 2019 (1), generating newer variants with higher infectivity (2). Delta (B.1.617.2), a dominant variant of concern (VOC) with higher severity (3, 4), had successfully outgrown the other variants (5), and caused several breakthrough infections (6). The newest VOC, Omicron, has caused major waves of infection across the world, but with significantly lower severity than Delta (7, 8). Before Delta, Alpha (B.1.1.7), another VOC, had higher transmissibility than its contemporary variants (3). Though variants such as Beta (B.1.351), and Gamma (P.1) were considered as potential VOCs at one point in time, they failed to dominate across the world. Immune evasion by the new variants against the antibodies generated against the previous variants or vaccines is natural during viral evolution and has been the case for Delta (9, 10). Though a trade-off between the virulence and transmissibility has been evident in several viral infections, there are exceptions as well (11). It is unclear if the subsequent SARS-CoV-2 variants have been adapting in humans causing more benign infections.

It is now fairly understood that the humoral immune escape coupled with increased transmissibility are important factors for a particular variant to gain dominance in the pandemic (12). Increased transmissibility is rendered by a number of factors including enhanced entry and better survival. Epithelial cells in the respiratory and intestinal systems are permissive to SARS-CoV-2 both *in vitro* and *in vivo* (13).

Innate immune response instructs the adaptive response through cytokines, chemokines, and antigen presentation (14). By far, there is no conclusive evidence of a productive infection of immune cells by SARS-CoV-2 (15, 16). Cytokine storm that has been implicated in the severe COVID-19 symptoms (17, 18) is an outcome of excessive secretion of pro-inflammatory cytokines first secreted by the epithelial cells and, in response by DC and other immune cells. Since viremia is not prominent in COVID-19 unlike in blood-borne viral diseases (19), the importance of the infected epithelial cells in triggering the adaptive responses is significant.

RLR pathway constitutes an important network recognizing double stranded RNA (dsRNA) intermediates of RNA viruses (20). Both RLR and TLR pathways are significantly impaired or delayed in COVID-19 patients (21–23) and validated in epithelial culture models (24, 25), contributing to COVID pathogenicity (18). Production of type-I IFNs and subsequent activation of JAK-STAT pathway are targeted by viral proteins and Alpha variant has evolved better mechanisms to evade innate response (25, 26). With a hypothesis that the newer and more successful variants are better in suppressing the innate responses, we investigated the details of RLR pathway activation in response to five different variants of SARS-CoV-2, including Delta. Our results demonstrate a steady progression in the capabilities of the subsequent variants over their previous ones in either delaying or efficiently suppressing innate immune response. Delta suppressed the host response pathways RLR-IFN and JAK-STAT most successfully and also resisted IFN treatment. Gene expression analysis uncovered that Delta suppressed host response in general including all major innate immune response pathways much more profoundly than Alpha, which itself was evidently more advanced than the previous variants. These suggested that Delta has been able to replicate in the host without alerting the innate signal pathways and this could possibly have resulted in delayed activation of adaptive response. Our findings could be important in the ever-changing contexts of COVID-19 symptoms and intervention strategies in addition to providing important clues to the evolutionary dynamics of SARS-CoV-2.

## RESULTS

### Delta genomic RNA has high replicative fitness in culture, but generates low infectious viral titers

Since Delta and Alpha variants had higher infectivity in the populations, we decided to compare their replicative and infectious fitness with the earlier variant isolates in time-course experiments in Caco2 cells. Previous studies have demonstrated that Caco2 cells are highly permissive to SARS-CoV-2 (13). In a comparative analysis, both lung epithelial cell line Calu3 and Caco2 showed comparable permissivity to SARS-CoV-2 (Supplementary Figures 1 A and B). Further, IRF3 phosphorylation at S396 residue in response to 1 MOI of SARS-CoV-2 infection was evident in Caco2 (as shown in the upcoming section), but not in Calu3 (Supplementary Figure 2), thus suggesting better suitability of Caco2 culture in our studies described in the following sections. S396 phosphorylation has been demonstrated to promote IRF3 nuclear translocation (27). Colon epithelium is a target of SARS-CoV-2, and intestinal distress being a major symptom in COVID-19, the choice of Caco2 is relevant to this study. Cells were infected with 1 MOI of five different SARS-CoV-2 variant isolates (B.6, B.1.1.8, B.1.36.29, B.1.1.7 (Alpha), and B.1.617.2 (Delta)) for up to 72 hpi, and the cellular and supernatant viral RNA titers and infectious titers were measured. Genetic variation among these variants has been depicted in Figure 1A. B.6 is an isolate of A3i clade that was prominent during the early part of the pandemic while B.1.1.8 belongs to A2a clade that diverged with a characteristic D614G conversion in Spike (S). B.1.36.29, another isolate of A2a clade, several cases of which was reported in India, has additional characteristic N440K mutation in the RBD of S. Intracellular RNA analysis revealed that Delta replicated most efficiently right from 24 to 72 hpi (Figure 1B), followed by Alpha, B.1.36.29, B.1.1.8, and B.6 in that order. Viral RNA levels in the supernatant followed similar trend (Figure 1C). However, the infectious titer data differed from the replication data where Delta displayed the least titers with B.1.36.29, and Alpha attaining the highest titers followed by B.1.1.8, and B.6 (Figure 1D). Thus, the higher rate of RNA replication of Delta did not translate into high infectious fitness. The relatively lower infectious titers of Delta also suggested that viral load may not be a major factor behind its higher transmissivity. Interestingly, spike (S) and nucleocapsid (N) immunoblots revealed that Alpha, and B.1.36.29 follow a pattern of high levels of S and N (Figure 1E) that correlated with their infectious titers, indicating that the higher availability of the structural proteins could be a determining factor in their higher infectious titers.

**Figure 1.**
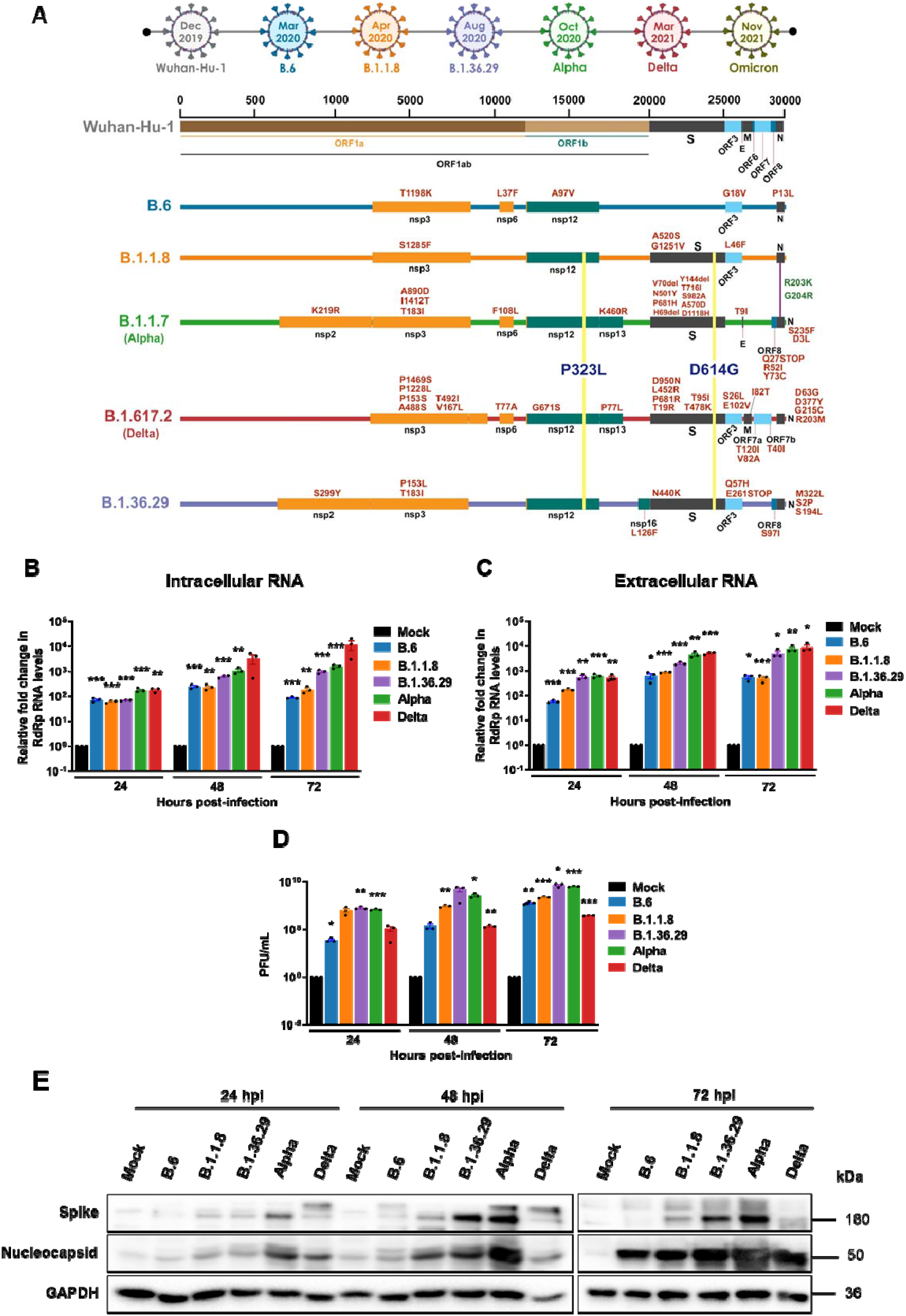
Delta has the highest RNA replication efficiency, but also has low infectious titers. (A) Schematic representing the mutations found in the five distinct variant isolates compared with the ancestral Wuhan isolate. The timescale on the top represents the month of the first reporting of the variant in GISAID. (B) Intracellular SARS-CoV-2 RNA quantified by real-time qRT-PCR detection of viral RdRp region. Caco2 cells were infected with 1 MOI of one of the five distinct variant isolates and incubated for the specific time-intervals as shown in the graph. The cells were harvested, RNA prepared from which SARS-CoV-2 RNA were analyzed by real-time qRT-PCR. The fold-changes against the mock-infected samples were generated through ΔΔ-Ct method by normalizing against the internal control *RNase P* of the corresponding sample. and have been plotted in the graph. (C) SARS-CoV-2 RNA in the culture supernatants quantified by real-time qRT-PCR detection of viral RdRp region. As in 1B, the fold-changes in the levels were plotted against the mock-infected samples for individual time points and normalized against *RNase P* values. (D) Infectious titers of SARS-CoV-2 from culture supernatants infected with the distinct variant isolate, determined by PFA. The culture supernatants collected at specific time-interval post-infection were cleared of the debris and were serially diluted and used as inoculum to infect fresh monolayers of Vero cells. The infected wells were layered with Agarose and the plaques formed were identified by staining with crystal violet. All the graphs contain results from biological triplicates. (E) Immunoblots detecting the levels of SARS-CoV-2 S and N proteins in the cells infected with the respective variant at specific time-interval. All graphs were prepared using GraphPad Prism version 8.0.2. Statistical significance is represented as *, **, and *** for p<0.05, p<0.01 and p<0.005 respectively.

### RLR and JAK-STAT are activated by early variants, but not by Delta

We next analyzed RLR mediated innate response to SARS-CoV-2 variants. IRF3 phosphorylation, a good measure of RLR activation, was activated by B.6 and B.1.1.8 variants by 48 hpi and continued until 72 hpi in Caco2 cells (Figure 2A). Though substantially delayed as reported elsewhere, the definite activation clearly suggested that the cells are able to overcome the suppression imposed early on by the virus. However, Alpha, Delta, and B.1.36.29 successfully inhibited IRF3 phosphorylation throughout 72 hpi, indicating that they have employed additional mechanisms to completely silence RLR activation. *IFNB1* expression at 24 hpi was limited to B.6 infection (Figure 2B) whereas by 48 hpi, strong induction was also found in B.1.1.8. A modest induction was visible in B.1.36.29 infection at 48 hpi. Phenomenal induction of *IFNL1* by B.6 and B.1.1.8 right from 24 hpi and at moderate levels by Alpha indicated that it is regulated distinctly from *IFNB1* (Figure 2C). Intriguingly, Delta caused considerable induction of *IFNL1* at 24 hpi, that faded progressively with time. The induction of *IFNB1* and *IFNL1* in the absence of IRF3 phosphorylation suggested that they are activated by IRF3-independent mechanisms during SARS-CoV-2 infection. B.1.36.29 most successfully suppressed both *IFNB1* and *IFNL1* activation. STAT1 phosphorylation in B.6 and B.1.1.8 infections by 24 hpi confirmed IFN-mediated activation of JAK-STAT pathway (Figures 2 A and D). Alpha infection delayed STAT1 phosphorylation till 72 hpi while Delta induced a modest and steady phosphorylation since 24 hpi. IFIT1 and its transcript levels closely mirrored STAT1 phosphorylation (Figures 2 A, E, and F). Similar inductions of MDA5 and its transcript *IFIH1* (Figure 2 G and H respectively), and *DDX58* transcripts (Figure 2I) further confirmed a strong activation of ISGs in B.6 and B.1.1.8 infections, and modest induction in Alpha, but insignificant in Delta and B.1.36.29 infections. These results indicated that the earlier variants indeed caused delayed RLR activation, but the later variants Alpha and Delta, in that order, progressively gained better mechanisms to effectively silence it. It is intriguing though that B.1.36.29 that suppressed RLR response more efficiently than Alpha had emerged well before it, but could not become a dominant variant.

**Figure 2.**
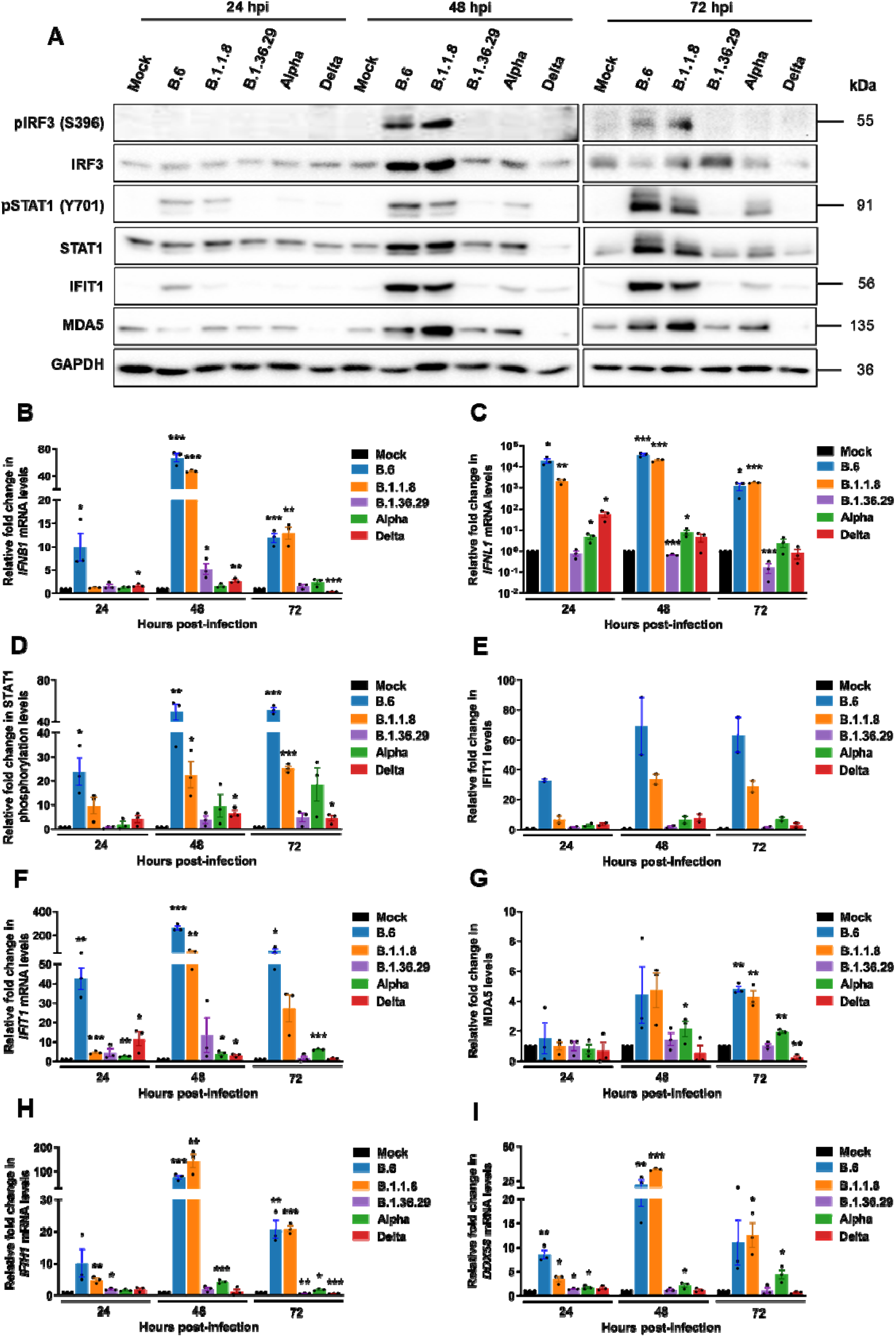
Delta infection causes long-term and complete suppression of RLR and JAK-STAT pathways. (A) Immunoblot images demonstrating the phosphorylation of IRF3 and STAT1 along with the expression of ISGs IFIT1 and MDA5 in Caco2 cells infected separately with one of the five variant isolates. (B) qRT-PCR quantification of *IFNB1* transcripts in cells infected with the individual variants. (C) Similar quantification for *IFNL1* transcripts. (D) Densitometric quantification of the phosphorylation of STAT1 across the infected samples. (E and F) Densitometric quantification of IFIT1 and qRT-PCR analysis of its transcripts respectively (n=2). (G) Densitometric quantification of MDA5 expression and (H) qRT-PCR analysis of its transcripts. (I) qRT-PCR quantification of *DDX58* transcripts in individual infections. All the graphs are representatives of biological triplicates. All graphs were prepared using GraphPad Prism version 8.0.2. *GAPDH* was used as the normalization control for qRT-PCR. Statistical significance is represented as *, **, and *** for p<0.05, p<0.01 and p<0.005 respectively.

### Delta and Alpha evade IFN response, but are partially susceptible to RLR activation by poly (I:C)

Our results clearly demonstrated that B.6, B.1.1.8 and Alpha have progressively developed capabilities to delay RLR and IFN signaling pathways whereas Delta, and B.1.36.29 are further evolved to silence the responses throughout the infection time-course. The activation of RLRs by their ligands is dependent on their post-translational modification (28) and hence could be a potential target for suppression by Delta. We asked if Delta could evade the prior activation of RLR pathway where previously activated RLR pathway would be suppressed by its infection. Caco2 cells transfected with poly (I:C) for 12 h were infected with the variants for 24 h (Figure 3A). We have described in the earlier section that 24 h infection caused a moderate increase in STAT1 phosphorylation in B.6 and B.1.1.8 infections (Figures 2 A and D) and hence this timeframe would be ideal to study the impact of RLR activation. While the induction of *IFNB1* confirmed the activation of RLR following poly (I:C) (Figure 3C), treatment, STAT1 phosphorylation (Figures 3 B and D) accompanied by elevated IFIT1 and MDA5 levels (Figures 3 B, E, and F) indicated the activation of JAK-STAT pathways. Though poly (I:C) augmented STAT1 phosphorylation in B.6 infection, its extent was masked by the higher basal level of phosphorylation caused by the infection (Figures 3 B and D). A similar masking was also seen in *IFNB1* levels in B.6 infection (Figure 3C) that caused robust *IFNB1* activation at 24 hpi (Figure 2B). Interestingly, STAT1 phosphorylation at 36 h post-poly (I:C) transfection in the mock-infected samples was comparable with the those that were similarly transfected and infected by B.6 (Figure 3B, lanes 3 and 5 respectively), suggesting that poly (I:C) transfection resulted in the saturation of STAT1 phosphorylation. The treatment resulted in appreciable drop in N levels in B.6, B.1.1.8, and B.1.36.29 infections, but not in Alpha and Delta variants (Figures 3 B). Poly (I:C) inhibited RNA replication of all variants (Figure 3G), indicating that genomic RNA replication of all SARS-CoV-2 variants are susceptible to the prior activation of RLR pathway.

**Figure 3.**
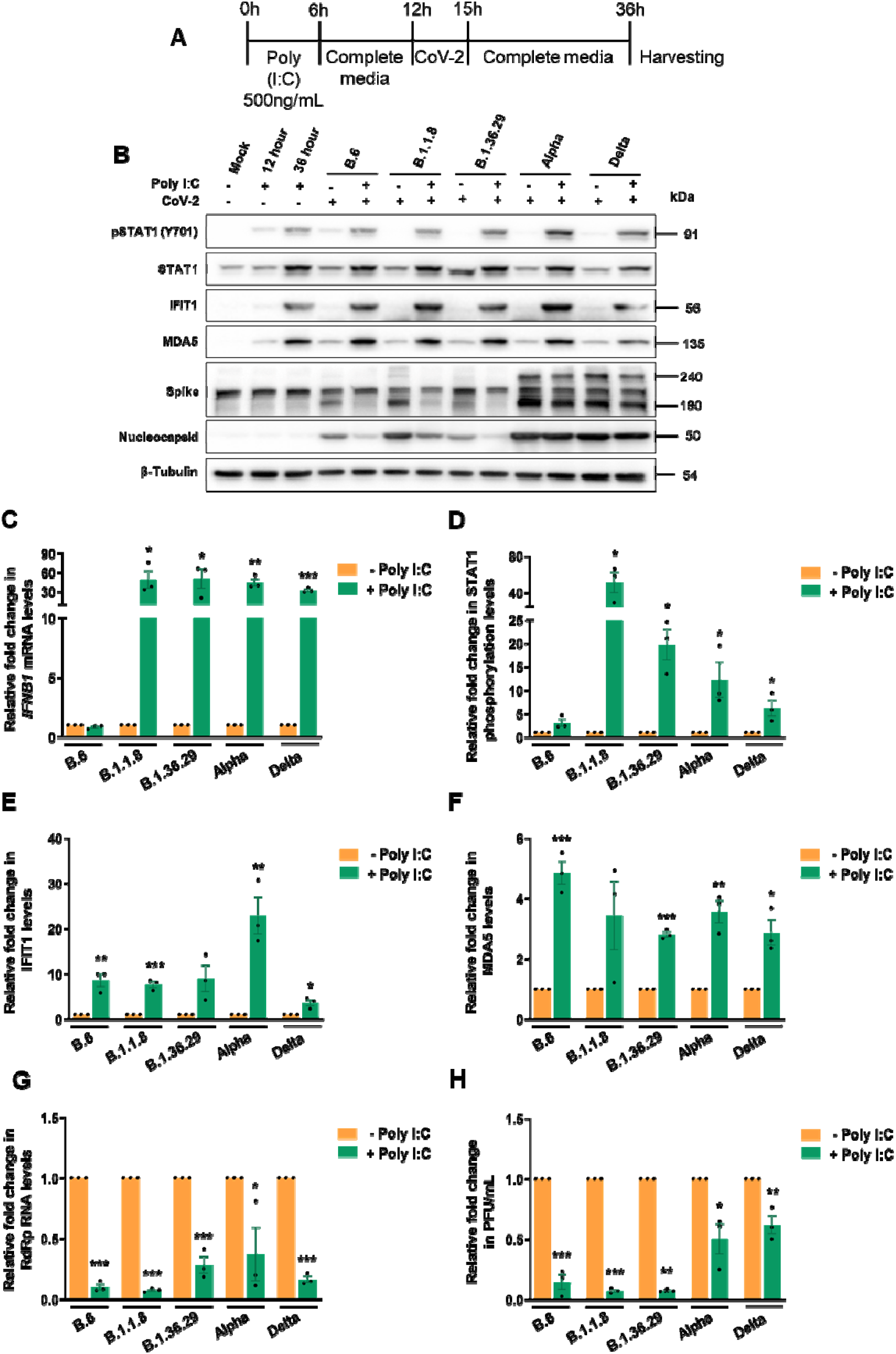
Alpha and Delta are sensitive to RLR activation by poly (I:C). (A) Schematic of the experimental set up for poly(I:C) treatment prior to variant infections. Caco2 cells were transfected with 500 ng/mL poly (I:C) for 6 h after which the transfection media replaced with growth media for incubation for another 6 h. At this point, the cultures were infected with 1 MOI of individual variant with 3 h of inoculation followed by further incubation in virus-free medium for a total of 24 h infection. (B) Immunoblots of the samples prepared from the infection for analyzing JAK-STAT activation. (C) *IFNB1* quantification in poly(I:C) treated, infected samples against the untreated, infected samples by qTR-PCR. The values are represented as fold-changes. First, fold-changes from the infected samples against the mock-infected samples were generated through ΔΔ-Ct method by normalizing against the internal control *GAPDH* of the the corresponding sample. Subsequently, fold-changes of such values generated in the poly (I:C) treated, infected samples against the untreated samples were calculated and plotted in the graph. (D-F) Densitometric quantification of the STAT1 phosphorylation and expressions of IFIT1, and MDA5. (G) SARS-CoV-2 RNA levels in the supernatants of poly (I:C) treated, infected samples measured by qRT-PCR and represented as fold-changes against the values from the respective untreated, infected samples. First, fold-changes from the infected samples against the mock-infected samples were generated through ΔΔ-Ct method by normalizing against the internal control *RNase P* of the the corresponding sample. Subsequently, fold-changes of such values generated in the poly (I:C) treated, infected samples against the untreated samples were calculated and plotted in the graph. (H) Infectious titers of SARS-CoV-2 in the supernatant of poly (I:C) treated, infected samples measured by PFA. All the graphs are representatives of biological triplicates. All graphs were prepared using GraphPad Prism version 8.0.2. Statistical significance is represented as *, **, and *** for p<0.05, p<0.01 and p<0.005 respectively.

However, poly (I:C) had only partial impact on the infectious titers of Alpha and Delta while the other variants were susceptible (Figure 3H). These results clearly indicated that early activation of RLR pathway prior to infection is efficient enough to restrict SARS-CoV-2 but the later variants are able to partially overcome this restriction. We then studied the sensitivity of SARS-CoV-2 variants to type-I IFN. Though SARS-CoV-2 proteins are shown to intercept STAT1 phosphorylation leading to its inactivation, IFNs are also shown to restrict SARS-CoV-2 replication (24, 29). IFN-α treatment of Caco2 cells (Figure 4A) activated JAK-STAT pathway, evident from increased STAT1 phosphorylation (Figures 4 B and C) and elevated levels of IFIT1 and MDA5 (Figures 4 B, D, and E). The treatment brought about considerable reduction in N levels in B.6, B.1.1.8, and B.1.36.29, but not in Alpha and Delta infections (Figures 4 B). IFN-α treatment caused significant drop in viral RNA titers in B.6, B.1.1.8, and B.1.36.29, but much less for Alpha and Delta infections, with Delta displaying the highest resistance (Figure 4F). Infectious titers of B.6, B.1.1.8, and B.1.36.29 were also significantly lower upon IFN-α treatment, but not much of Alpha and Delta (Figure 4G), indicating that the latter two variants have acquired resistance to IFN-α signaling, but are susceptible to RLR pathway activation. These results also suggest that the poly (I:C)-mediated restriction of SARS-CoV-2 is less dependent on IFN pathways, but uses non-canonical mechanisms against which SARS-CoV-2 has not gained resistance. Collectively our results indicated a gradual and independent evolution of mechanisms to resist IFN-dependent and independent antiviral mechanisms by the recent SARS-CoV-2 variants.

**Figure 4.**
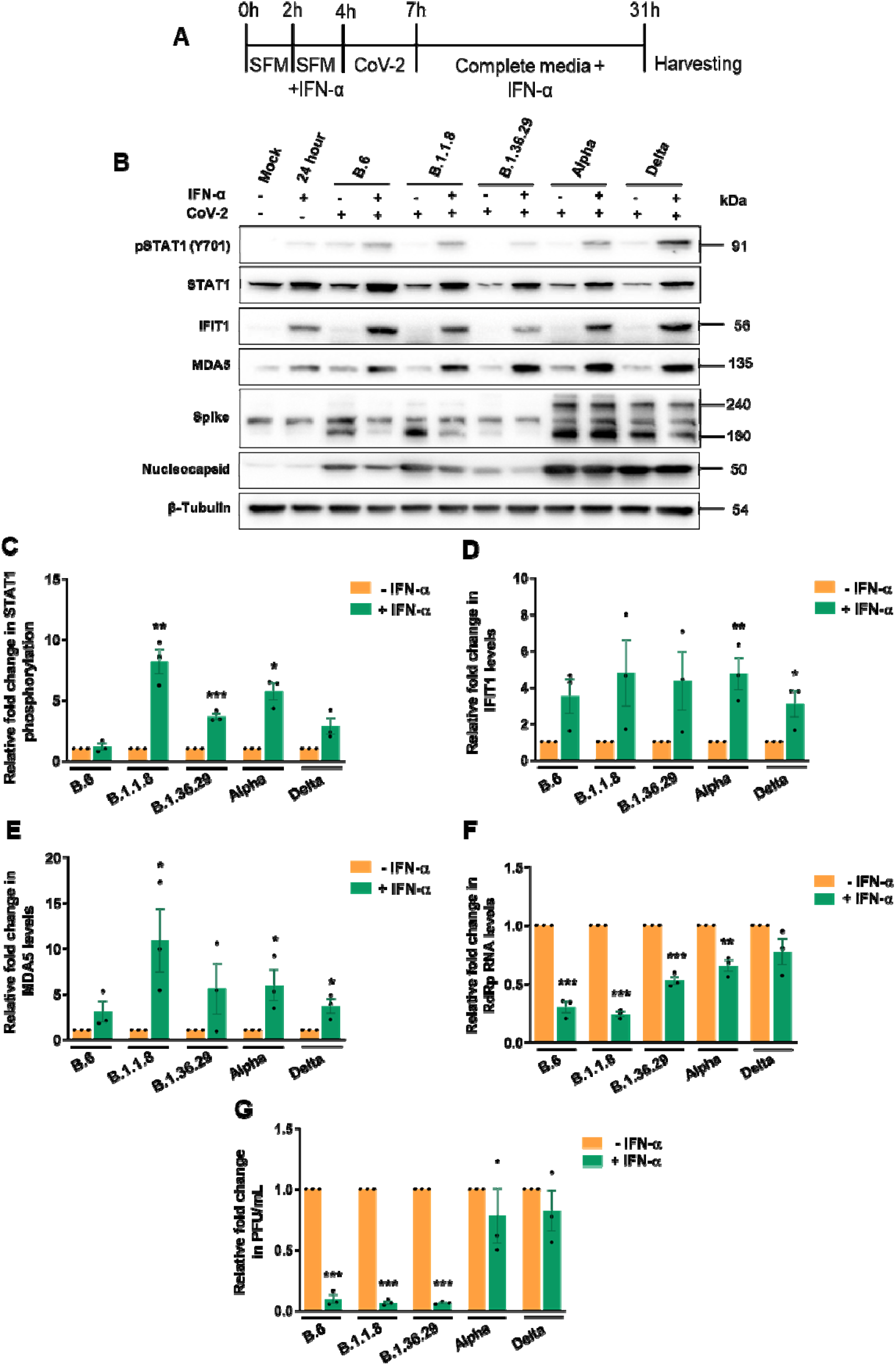
Alpha and Delta show resistance to IFN. (A) Schematic of the experimental set up for IFN-α treatment prior to variant infections. 2 h prior to IFN-α treatment, Caco2 cells were incubated with SFD. IFN-α containing SFD was added to the cells at a concentration of 500 U/mL of IFN-α and further incubated for 4 h. Cells were infected in SFD in the absence of IFN-α for 3 h after which the inoculum was replaced with grown media containing IFN-α and incubated until 24 hpi. (B) Analysis of the JAK-STAT pathway activation following IFN-α treatment by immunoblotting STAT1 phosphorylation and expressions of IFIT1 and MDA5. (C-E) Measurement of STAT1 phosphorylation and expressions of IFIT1, and MDA5 by densitometry. (F) Measurement of SARS-CoV-2 RNA levels in the supernatants of IFN-α treated, infected samples by qRT-PCR, which is represented as fold-changes against the values from the respective untreated, infected samples. As in Figure 3G, fold-changes from the infected samples against the mock-infected samples were generated first through ΔΔ-Ct method by normalizing against the internal control *RNase P* of the corresponding sample. Subsequently, fold-changes of such values generated in the poly (I:C) treated, infected samples against the untreated samples were calculated and plotted in the graph. (G) Infectious titers of SARS-CoV-2 in the supernatant of IFN-α treated, infected samples measured by PFA. All graphs were prepared using GraphPad Prism version 8.0.2. Statistical significance is represented as *, **, and *** for p<0.05, p<0.01 and p<0.005 respectively.

### Gene expression profiling reveals strong inactivation of antiviral pathways by Delta

We analyzed the time-course transcriptional reprograming (TR) following infections by individual variants of SARS-CoV-2 except B.1.36.29 in Caco2 cultures. B.1.36.29 infection was not included as this variant lacked the advanced feature of IFN resistance and was not a prominent variant in circulation. Principal component analysis (PCA) confirmed that the biological replicates clustered together and maximum variance was observed for B.6 and B.1.1.8 followed by Alpha from the controls while Delta showed the least variations (Figure 5A). This suggests strong host transcriptional response to B.6, B.1.1.8 and Alpha, but not to Delta. We further performed differential expression analysis for the four variants against control, to identify significantly regulated genes (FDR < 0.05 and absolute log2 fold change > 1). The number of differentially expressed genes (DEGs) suggests that B.6 caused the sharpest response followed by B.1.1.8 and Alpha in that order (Figure 5B). Alpha caused a comparable scale of TR at 72 hpi, but was significantly delayed compared to B.6 and B.1.1.8, indicating a better control of host response by this variant. Since IRF3 phosphorylation remained muted, and *IFNB1* and *IFNL1* levels were uninduced even at 72 hpi by Alpha (Figures 2A-C), this late surge of host response is likely to have been coordinated by IFN-independent mechanisms. Unlike B.6, B.1.1.8, and Alpha infections, Delta caused steady, benign TR throughout the time-course, suggesting that these variants have been able to effectively contain multiple surveillance mechanisms of the host and thus have a stricter control over host responses (Figures 5B). This trend of progressive delay in the host responses to B.1.1.8, and Alpha, and the mild response to Delta indicated that the lately emerged variants have better mechanisms to evade the host surveillance than their earlier variants. Interestingly, the overall distribution of DEGs fold-change by Alpha remained much lower than those from B.6 and B.1.1.8 (Figure 5C). Highest distribution for B.6 and B.1.1.8 was found at 48 hpi while for Alpha, it was seen at 72 hpi, confirming that Alpha has evolved to delay the innate immune response, probably not to evade it totally. Unlike in the case of other variants, the distribution of DEGs fold-change was maintained throughout the time-course in Delta infection, suggesting that it is able to tightly suppress the host response. Analysis of the consolidated DEGs for the variants indicates that the TR imprint of Alpha resembled more with those of B.6 and B.1.1.8 than it did with Delta, while that of Delta overlapped closely with both B.6 and Alpha (Figure 5D). GO analysis of the consolidated DEGs demonstrated a strong enrichment of genes participating in antiviral response for the up-regulated genes in B.6, B.1.18, and Alpha infections, but not in Delta (Figure 5E). Further, mononuclear differentiation, and leukocyte migration factors were strongly enriched in B.6, B.1.18, and Alpha infections, as compared to Delta, indicating that Delta infection does not alarm the adaptive immune response (Figures 5E), particularly from 48 hpi (Supplementary Figure 3A). Stronger enrichment of DEGs from 48 hpi underlined the delayed response to SARS-CoV-2. KEGG analysis identified substantially reduced enrichment of genes involved in cytokine-chemokine, NF-κ B, TNF, NLR, and PI3K-AKT signaling pathways in Delta infection as compared with B.6, B.1.1.8 and Alpha infections (Figures 5F, and Supplementary Figure 3B). Among the down-regulated genes in B.6, B.1.1.8, and Alpha infections, enrichment was found for processes involved in fatty acid metabolism, and lipid localization particularly beyond 48 hpi, indicating unique associations of Delta with the host-derived membranous compartment (Figures 5G, and Supplementary Figure 4A). Membrane components being very critical for SARS-CoV-2 life-cycle, their metabolism is modulated by the viruses for their benefit. Interestingly, nucleotide and alcohol metabolism were also down-regulated by these variants. Though B.6 and Delta infections caused transcriptional downregulation of a number of genes at 24 hpi (Figure 5C), no significant functional enrichment was observed for these genes from Delta samples (Supplementary Figures 4 A and B). KEGG enrichment analysis showed a Delta-specific downregulation of a small set of components of pro-inflammatory IL-17 and TNF signaling pathways, and cytokine-cytokine interaction, late in infection (Figure 5H, and Supplementary Figure 4B), indicating that Delta not just spares cytokine induction, but inhibits it at the later stages of infection. The progressively depleting proportion of the regulated genes shared by B.6 with B.1.1.8, Alpha, and Delta in that order indicated a continuing divergence of the evolving variants from the earliest variant B.6 (Supplementary Figure 5A). Only a small fraction of DEGs across all time-points overlapped among the four infections to form a common pool of commonly regulated genes (261 up- and 57 down-regulated), indicating the unique transcription profiles generated by the individual variants (Supplementary Figures 5A, 6A, and 6B). Among the 261 commonly up-regulated genes, significant enrichment was seen for antiviral response processes in GO analyses (Supplementary Figure 5B). Lack of enrichment for genes uniquely associated with individual variants indicated that the functional significance of a significant proportion of DEGs cannot be ascertained for each of the variants (Supplementary Figures 5 B and D). The commonly down-regulated genes (Supplementary Figure 5D) did not form any enrichment while those from B.6, B.1.1.8, and Delta formed individual enrichment groups (Supplementary Figures 5 C and E). Delta caused the lowest magnitudes of gene activation and suppression among the common set of DEGs across the variants. (Supplementary Figures 6 A and B).

**Figure 5.**
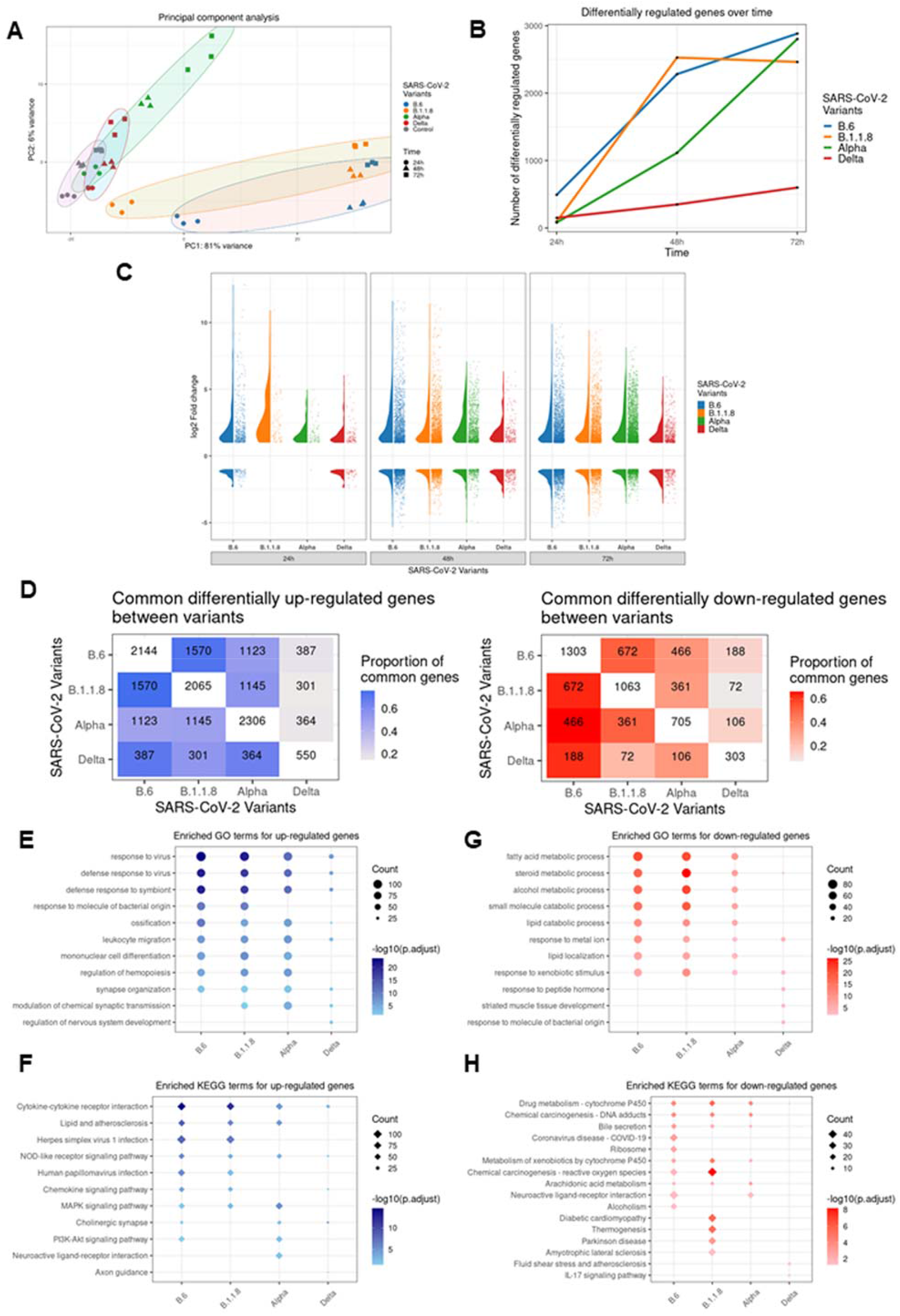
Gene expression profiling in response to SARS-CoV-2 variants. Total RNA isolated from Caco2 cultures infected with distinct variants for the respective time-intervals were subjected to next-generation sequencing. Three biological replicates were used for library generation and sequencing. The sequences generated were analyzed by PCA. All DEGs considered had Log2 fold-changes >1 for the up-regulated and <-1 for the down-regulated genes, with *p adjacent* value <0.05. (A) PCA analysis of the sequences generated. Regularized log transformed count data was used for computing principal components. The PCA confirms the quality of data where biological replicates clustered together. From the control samples, maximum variance was observed for B.6, B.1.1.8 followed by Alpha (B) Line graphs showing the total number of differentially expressed genes in response to individual variant at the specified time points. The number represents sum of both up- and down-regulated genes. (C) Violin plots representing the distribution of log2-fold changes and jitter plot representing number of DEGs in response to different variants at each time-points. (D) Heat-maps representing the overlapping DEGs across time-points between variant-infected samples in x-axis and y-axis. The numbers in diagonal boxes represent the total number of statistically significant up- or down-regulated genes in the corresponding samples. The up-regulated genes are represented in blue boxes while the down-regulated ones are in red. The color intensity represents the proportion of DEGs for the variants in y-axis overlapping with DEGs for the variants in the x-axis. (E and F) Enrichment analysis representing the Enriched GO (circles) and KEGG (diamond) terms for up-regulated DEGs for each variant-infected sample across time-points. Size of the dot is proportionate to the number of DEGs representing the enriched term and the intensity of the color represents the -log10 (adjusted p-value) of the DEGs represented. (G and H) Similar enrichment analysis for down regulated genes caused by infection by individual variants. The up-regulated DEGs are represented in blue color while the down-regulated ones are in red.

### Delta infection causes more intense and persistent subversion of cytokines, chemokines, and antigen presentation genes than Alpha

Delta not only caused a low-grade TR of antiviral genes, but lower quantum as well (Figure 6 A and B) maintaining a steady profile with no major changes during the time-course, further suggesting that they have developed capabilities to persistently silence the response. The violin plot considered 822 genes classified under various processes contributing to innate immune response, response to cytokine, defense response, type-I IFN pathway, and leukocyte activation and differentiation. In line with our earlier data, only a small set of genes were reprogrammed by Delta infection (Figures 6A). Within a select subset of these genes, Delta specifically down-regulated several antiviral genes of interest such as *OASL*, *NLRC5*, *IFNL2* and *IFNL3* at 72 hpi (Figure 6B). Down-regulated genes in Delta also enriched for cytokine receptor interaction, TNF and IL-17 signaling (Figure 6B and Supplementary Figure 4B), indicating the distinct influence of this variant on the host response. Alpha and Delta suppressed type-I IFN induction whereas B.6 and B.1.1.8 induced *IFNB1* right from 24 hpi. Type-III IFNs, the early responding cytokines in epithelial cells, were detected early in B.6 and B.1.1.8 infections and later in Alpha infection (*IFNL1, IFNL2* and *IFNL3*) at 48 hpi, thereby indicating that the late surge of TR in Alpha infection could partly be triggered by this class of IFNs (*IFNL2* and *IFNL3* in Figure 6B; *IFNL1* in Supplementary Figure 7). Consistent with these observations, ISG activation was also very limited in Delta infection (Supplementary Figures 7 and 8A). RLR and NLR pathway components were also significantly activated by B.6, and B.1.1.8, and to a moderate level by Alpha (Supplementary Figures 8B and 9A respectively). Absence of any appreciable activation of NF-κB by Delta as compared with the others was in agreement with the earlier observations (Supplementary Figure 9B). Intriguingly, despite a clear absence of both type-I and – III IFNs, a limited set of ISGs (*OAS2* and a few *IFIT*s) were activated by Delta indicating the activation IFN-independent pathways (Figure 6B and Supplementary Figure 7). Pro-inflammatory chemokines *CCL4* and *IL-6* that promote cytokine storm were activated only by B6 and B.1.1.8 (Figure 6B and Supplementary Figures 8 and 10), indicating that the magnitude of cytokine storm in Delta infections could be much smaller than that by the earlier variants. However, TNF-α expression was detected in Delta infection (Figure 6B, Supplementary Figures 9B and 10C), albeit late and milder than the previous variants, indicating that its regulation is independent from that of *CCL4* and *IL-6*. These data, agreeing with the immunoblot data (Figures 1 and 2) confirm that the lately emerged variants have evolved mechanisms to suppress both type-I and –III IFN, as well as cytokine and chemokine activations. Additionally, antigen presentation was also compromised in Delta infection. A study had previously reported the inhibition of activation of MHC Class-I pathways by SARS-CoV-2 where they analyzed the results until 24 hpi (30). We detected similar results, but found their activation at later hours of infection. While the regulators *NLRC5*, *IRF1* and *STAT1*, and *HLA-B* were progressively activated through the infection time-course in B.6 and B.1.1.8 infections, they were hardly detected in Delta infection (Figure 6C, and Supplementary Figures 7E and 10). Collectively, our results demonstrate that Delta infection causes very mild response from the host cells thereby possibly resulting in a delayed or milder activation of adaptive immune response.

**Figure 6.**
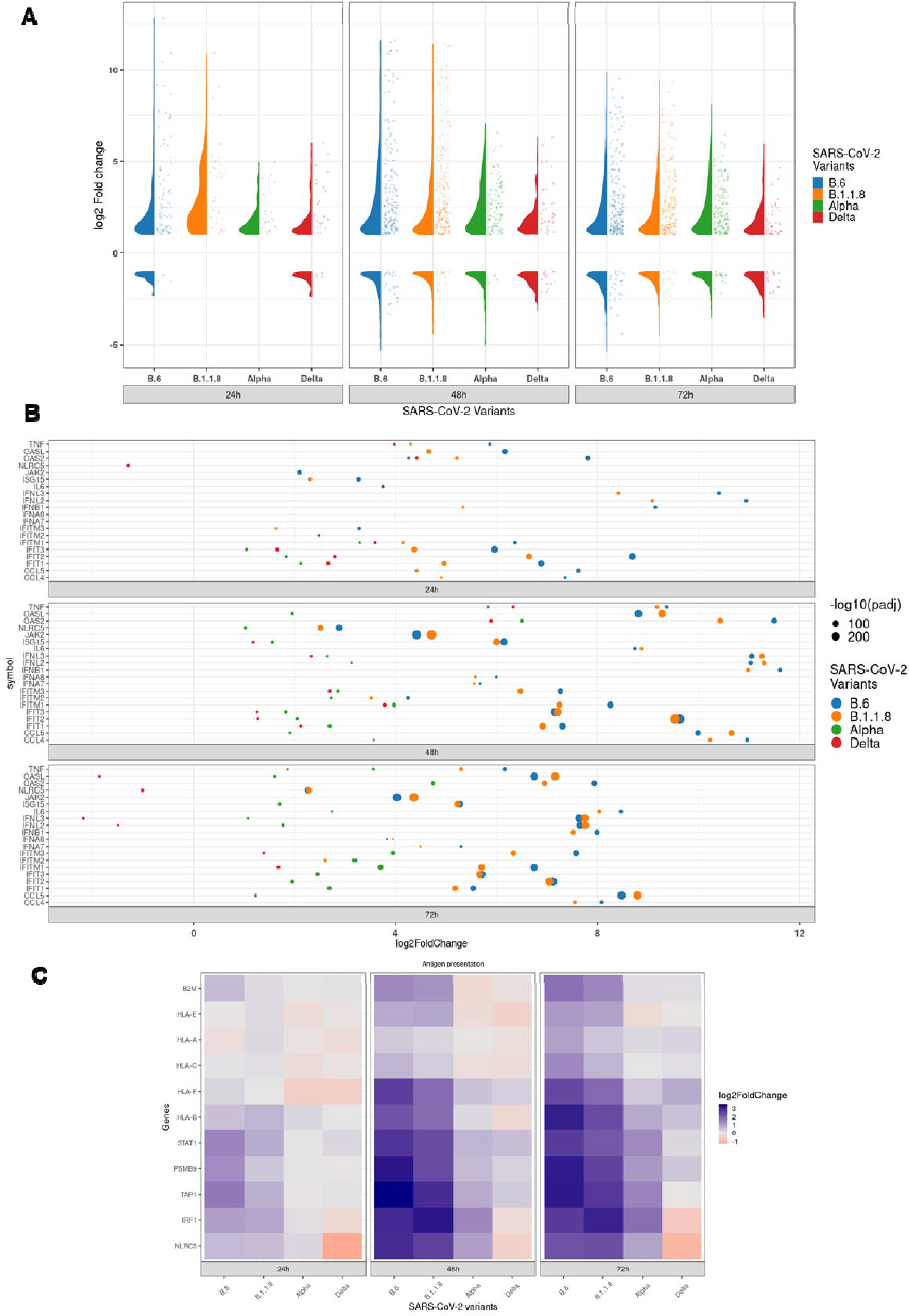
Activation of antiviral immune genes is severely suppressed in Delta infection. (A) Violin plot representing the distribution of log2 fold-change and dot plot representing number of differentially expressed genes annotated for antiviral functions for specified time points. (B) Jitter plot representing the log2 fold-change of 20 select genes participating in the innate immune response. Size of the dot represents the -log10 adjusted p-value. (C) Heat-maps representing log2 fold change of DEGs participating in antigen presentation in response to the individual variants at the specified time-points.

## DISCUSSION

Viral infections are studied from the perspective of virulence and transmissibility, which often share diffused borders. Studies on viral virulence have often been impeded with theoretical and empirical studies running in parallel. Recent developments in the sequence determination of variants have given better insight into the process of viral and virulence evolution (11). Traditional wisdom suggests that the virulence caused by a pathogen in a new host would be tempered over a period of their co-existence driven by natural selection. R_0_, the pathogen fitness index, is proportionate to the ratio of transmission rate and the sum of the mortality and recovery rates. Though a trade-off between the virulence and transmission rate is often observed during the evolution of the relationship with the host, it may not be necessary (11, 31). In this study, we attempted to comprehensively characterize how the new variants that emerged during the pandemic timescale have evolved with the host from the point of the host-response to these individual variants. Our study in SARS-CoV-2 infected cells clearly demonstrates a spectrum of host response triggered by distinct viral variants where the earliest one B.6 caused the quickest, while the latest one Delta caused the most benign response. The responses against the other two variants were indications of the measured progression of the virus to a more benign variant (Figure 7). The variants emerged later have evolved better mechanisms to delay and to silence the innate response than their previous ones, facilitating their longer stay in the infected host. By this criterion, they can be identified as more evolved. This trait is likely to improve with the newer dominant variants emerging after Delta, such as Omicron. It is evident that suppression of innate immunity and resistance to IFN were achieved through distinct mechanisms. Our findings have important implications on the therapeutic approaches involving IFN therapy against the emerging variants.

**Figure 7.**
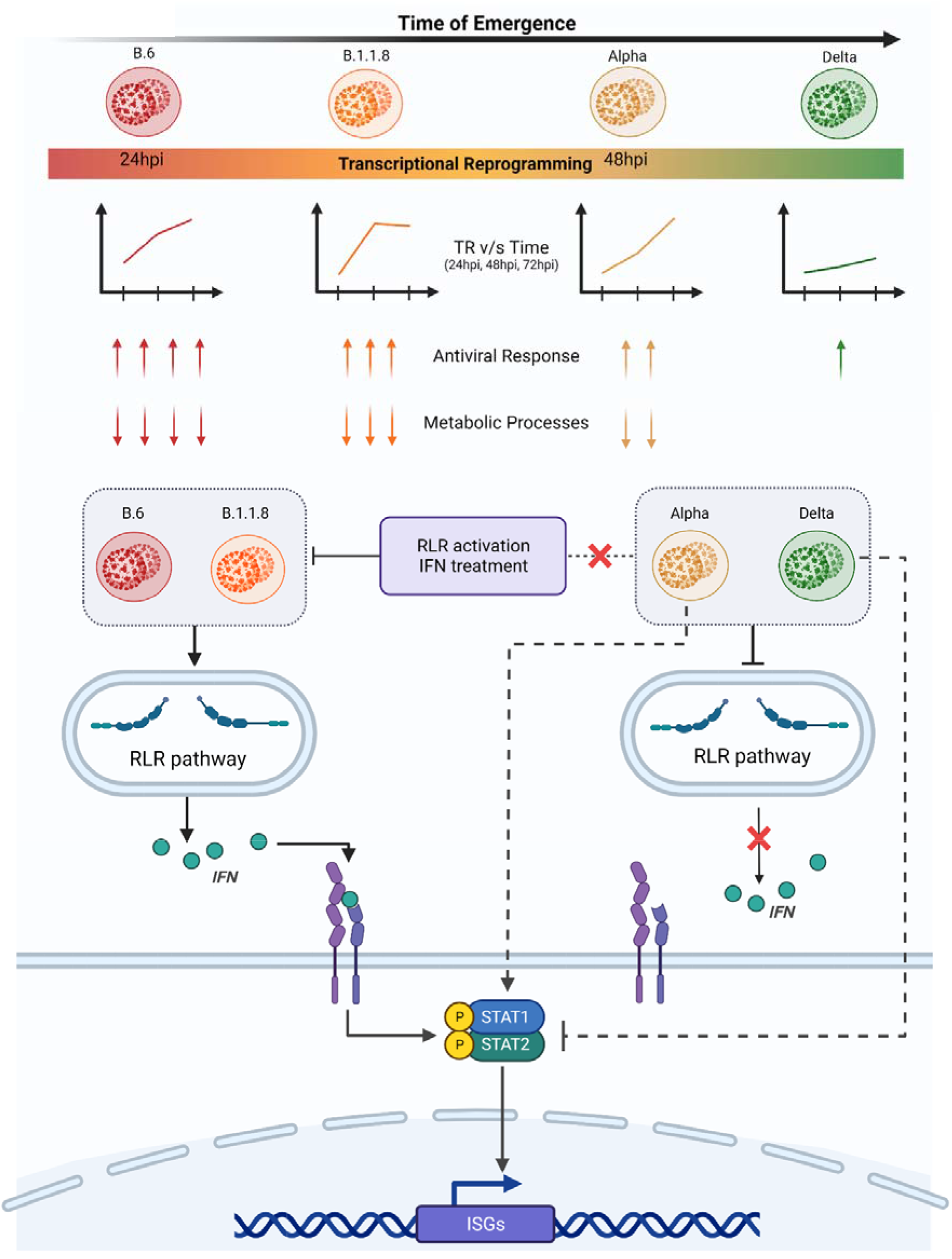
Delta variant has gained highly advanced control over the innate immune response and suppresses host responses effectively. The variants emerged during the early part of COVID-19 trigger moderate immune response by 24 h and robust response by 48 h post-infection. This was evident by the activation of RLR pathway that was further substantiated by transcriptome data. However, Alpha suppresses RLR pathway effectively, but failed to suppress STAT1 phosphorylation, possibly through IFN-independent mechanism. This was reflected in the late surge of transcriptional activities in Alpha infection. Delta has been the most advanced in suppressing not just innate immune response, but host response in general. Delta suppressed RLR pathway, IFN production and STAT1 phosphorylation, and this was reflected in the modest, steady response from the infected cells throughout the infection period. SARS-CoV-2 variants used in this study were presented based on their time of emergence from left to right with B.6 being the earliest and Delta being the most recent of them. The color of the variant virus particle shown in the schematic directly correlates with degree of transcriptional reprogramming by variants presented in graphical depiction below individual variants. The color intensity of the rectangular bar represents transcriptional reprogramming and control over host immune response by individual variant. Red represents elevated TR and strong activation of immune response while green represents lenient TR and greater control over host responses. Two of the GO enriched terms were presented with arrows. The numbers of arrows represent the potency of activation or inhibition, where up arrows indicate up-regulation of DEGs involved while down arrows indicate down-regulation. The variants studied here broadly fall under two groups based on the regulation of RLR pathway components and their response to activated innate immune responses (RLR activation by Poly I:C and JAK-STAT activation by IFN treatment). B.6 and B.1.1.8 activated RLR signalling followed by IFN secretion and ISGs expression via JAK-STAT axis. RLR and JAK-STAT signaling remain suppressed in Delta infection. Uniquely Alpha follows non-canonical mode of STAT activation without any detectable expression of IFNs.

Recent report on the evolution of Alpha to evade innate immune response more efficiently than its previous variants (25) was also captured in our studies as we did detect substantially elevated levels of N proteins in Alpha infections. Absence of overlapping mutations shared by Alpha, and Delta (Supplementary 11) indicates divergent mechanisms adopted by these variants to achieve similar outcomes. Clearly, they must have targeted the innate sensing pathways RLR and TLR uniquely. A much delayed, but strong host antiviral response against Alpha despite the induction of RLR pathway and *IFNB1* production indicated the involvement of alternate mechanisms that Delta was able to successfully suppress. In one such case, Delta was able to suppress the modest IFN-independent STAT1 phosphorylation and activation that was found in Alpha infections (Figures 2A, and 7). Minimal activation of *ISG15* by Alpha and Delta indicated that its suppression might be assisting these variants in lowering the ISGylation of its target molecules that are important mediators of innate immune response (Supplementary Figure 10G). Interestingly, despite a complete absence of IRF3 phosphorylation during Alpha infection, the host response exploded between 24-72 hpi, suggesting that this response is not orchestrated by IFNs.

Our studies also set a platform for further discussions on the larger question of the features that make a particular variant more transmissible. Though Delta RNA replication was the fastest in agreement with the existing literature (9), its infectious titers were lower than the previous variants, an indication that silencing the host response does not appear to provide it any particular advantage in terms of its viral load. However, silencing the innate response would be important from the point of view of the response of the host. With a reported higher R_0_ for Delta (32), there lacks a credible clinical data on its relative virulence compared with the previous variants in immunologically naïve populations. Delta indeed caused severe pathology during its emerging period while the majority population was unvaccinated. Currently, it appears highly improbable to conduct unbiased population studies on its severity due to the unavailability of immunologically naïve cohorts, as would the case of Omicron and future variants (33) . Based on our data, we could speculate that the contribution by the epithelial cells to the systemic responses could be significantly lower in persons infected by Delta as compared to its previous counterparts. A reduced communication from the epithelial cells would also result in lower adaptive response thus causing lower chances of cytokine storm. However, Delta-specific data on cytokine storm is lacking. At the same time, the absence of support from the adaptive immune response could result in persistent infection resulting in a more severe pathological damage in the respiratory and intestinal tissues. Long-term presence of active SARS-CoV-2 in patients is an indication of such a strategy (34). Whether this scenario contributed to higher cases of respiratory sickness associated with Delta infection needs further investigation. Higher incidences respiratory support and ICU admissions were reported in Delta prevalent regions, indicating that the lung could have been subject to more serious damage.

One potential criticism against our study could possibly be that these studies were not performed in human primary epithelial cells. However, the major objective of our study was to map the evolving trend of innate immune escape by the emerging variants and hence we needed a system that responds to the earlier variants. Animal models such as ACE2 transgenic mice and Syrian hamsters are also not natural hosts of SARS-CoV-2 and hence may not be a good choice to study the viral evolution as in the case of Influenza (35). Caco2, being colon epithelial cells of human origin and being highly permissive, have allowed us to study our objective thoroughly. The results from these studies could be of great significance in characterizing the ever-evolving nature of COVID-19.

## METHODS

### Cell culture, poly (I:C) transfection and IFN-α treatment

Vero (CCL-81) cells were purchased from Sigma-Aldrich cultured in complete Dulbecco’s modified eagle medium (cDMEM; Gibco) containing 10% Fetal bovine serum (FBS; Hyclone), and 1× penicillin-streptomycin cocktail (Gibco) at 37°C and 5% CO_2_. Caco2, purchased from ATCC, were grown similarly, but supplemented with 20% FBS. Cells were continuously passaged at 70-80% confluency and were maintained in a condition of ambient temperature and humidity.

Poly (I:C) transfections were performed as in previous report (36). Cells were seeded to reach 80% confluency. Transfection mix containing OptiMEM-Lipofectamine 3000-poly (I:C) was prepared according to manufacturer’s protocol and added to cells and incubated for 6 h. Later, the transfection mix was replaced with cDMEM and further incubated for 6 h. 12 h later, the transfected cells were infected with virus for 3 h and further incubated in fresh cDMEM until harvested for analyses.

For IFN-α treatment, cells were seeded to reach 80-85% confluency. Cells were supplemented with serum-free DMEM (SFD) for 2h for serum starvation. Later, SFD was replaced with fresh SFD containing 500U/mL IFN-α for 2h. Following this, the cells were infected for 3 h as earlier and incubated further with fresh cDMEM containing either PBS (vehicle) or 1000 U/mL (PBL Assay Science) of IFN-α and incubated for 24 h. Cells were harvested and used for RNA or protein work.

### SARS-CoV-2 isolates

Five variant isolates of SARS-CoV-2 used in this study were isolated (30) at the Centre for Cellular and Molecular Biology in the biosafety level-3 facility. Their genomes were sequenced (GISAID ID: EPI_ISL_458067; virus name-hCoV- 19/India/TG-CCMB-O2/2020 (B.6), EPI_ISL_458046; virus name-hCoV-19/India/TG-CCMB-L1021/2020 (B.1.1.8), GISAID ID: EPI_ISL_539744; virus name- hCoV-19/India/TG-CCMB-AC511/2020 (B.1.36.29), GISAID ID: EPI_ISL_1672391.2; virus name- hCoV-19/India/TG-CCMB-BB649-P1/2020 (B.1.1.7), GISAID ID: EPI_ISL_2775201; virus name- hCoV-19/India/TG-CCMB-CIA4413/2021 (Delta). The viruses were propagated in Vero (CCL-81) cells grown in SFD.

### Virus Infection, quantification, and titration

Caco2 cells were infected at 1 MOI for 3 h in SFD after which the inoculum was replaced with complete media and further grown until harvesting. Supernatants collected were processed for RNA preparation using Nucleospin Viral RNA isolation kit (Macherey-Nagel GmbH & Co. KG), and infectious titer assay (plaque formation assay (PFA)). qRT-PCR to quantify SARS-CoV-2 RNA was performed on Roche LightCycler 480 using nCOV-19 RT-PCR detection kit from Q-line Molecular. Infectious titers of the supernatants were calculated using PFA as mentioned previously (37). The viral supernatants were serially diluted to prepare inocula that were inoculated on cultured Vero cells. Post-infection the monolayers were immobilized by soft-agar medium and further incubated until the plaque were formed. Plaques formed from the replicates were counted, extrapolated to 1 mL volume by applying the dilution factor, averaged, and represented in plaque forming unit/mL (PFU/mL).

### Real-time quantitative RT-PCR

Cellular RNA samples were prepared using MN Nucleospin RNA kit (Takara). Equal quantities of RNA were reverse transcribed using Primescript Reverse transcriptase (Takara) following the manufacturer’s protocol. 50 ng of cDNA was used for quantification using SYBR Green mastermix (Takara) on Lightcycler 480 instrument (Roche). Transcripts of the host origin were normalized against GAPDH. Relative fold-changes between the experimental and control samples(2^ΔΔCt^) were calculated by represented in the graphs.

### Antibodies and immunoblotting

All primary antibodies were purchased from Cell Signaling Technologies except the anti-Spike antibody (Novus Biologicals), and anti-Nucleocapsid, anti-Tubulin and anti-GAPDH (Thermo Fisher) antibodies. HRP-conjugated secondary antibodies was purchased from Jackson ImmunoResearch. Protein pellets were lysed in an NP-40 lysis buffer as described earlier (36). Protein quantification was done using BCA method (G Biosciences). The immunoblots were developed on a BioRad Chemidoc MP system using ECL reagents (ThermoFisher and G Biosciences). Quantification was performed using ImageJ.

### Next generation sequencing

Library preparation was done using the MGIEasy RNA Library Prep Set (MGI) according to the manufacturer’s instructions. In brief, 500 ng total RNA was used as starting material from which ribosomal RNA was depleted using Ribo-Zero Plus rRNA Depletion Kit (Illumina). The rRNA depleted samples were fragmented, reverse transcribed and the second strands were synthesised. DNA was then purified using DNA Clean Beads provided in the kit followed by end repair and A-tailing. Barcoding and adaptor ligation were performed and the samples were purified. Samples were amplified using adaptor specific primers and quantified using Qubit dsDNA high sensitivity kit (Thermo Scientific). Sample fragment size was determined using 4200 Tape Station (Agilent). The samples were denatured and single stranded circular DNA strands were generated. Further, rolling cycle amplification was performed to generate DNA nanoballs. The samples were subsequently loaded onto the flow cells (FCL) and sequenced at PE100.

### Data Processing and Analysis

MGI adapters and low-quality reads were removed from raw sequencing reads using Cutadapt (38). Reads with quality scores less than 20 and smaller than 36 bp were discarded. The processed reads were then mapped to the human genome GRCh38 using Hisat2 with default parameters (39). Uniquely aligned reads were counted using feature Counts of Subread package (40). Count information was available for 60683 genes in the gtf file, downloaded from Ensemble (41). Genes with total 10 read counts across all the samples were removed resulting in 35906 genes for further analysis. Differential gene expression analysis was performed using DESeq2 (42). Genes with adjusted p-value < 0.05 and absolute log2 Fold change > 1 were considered differentially expressed. For PCA plot and heat map, the raw read counts were rlog normalized, available with the DESeq2 package.

### Functional enrichment analysis

Functional enrichment analysis was performed using clusterProfiler (43) for GO term and KEGG pathways enrichment. We only used the Biological process for GO term enrichment analysis. Similar enriched terms were further merged using the ‘simplify’ function of clusterProfiler with similarity cut-off set to 0.7. ‘p.adjust’ was used as a feature to select representative terms and ‘min’ was used to select features. ‘Wang’ was used as a method to measure similarity. Top 10 GO terms and KEGG pathways based count of genes were plotted.

### Statistical analysis

Statistical significance was calculated by paired end, two-tailed Student’s t-test method. All experiments were conducted minimum three independent rounds and averaged values are represented as scatter plot with bar graphs (depicting individual values of independent experiments). Error bars are representations of the mean ± SEM. All graphs were prepared using GraphPad Prism version 8.0.2. Statistical significance is represented as *, **, and *** for p<0.05, p<0.01 and p<0.005 respectively.

### Data availability

RNAseq data was deposited into GEO database under accession number GSE193122.

### Institutional biosafety

Institutional biosafety clearance was obtained for the experiments pertaining to SARS-CoV-2.

#### Institutional ethics clearance

Institutional ethics clearance (IEC-82/2020) was obtained for the patient sample processing for virus culture.

## Author contributions

D.T. and K.H.H. conceived and designed this study. D.T., V.S., S.P., and K.H.H. designed the experiments. D.T., V.S., S.P., D.G., and S.S. performed the experiments. N.K.S., D.T.S., and V.S. analysed the NGS data with assistance from D.T. K.H.H. wrote the manuscript with editing from D.T., and inputs from N.K.S. and D.T.S.

## Acknowledgement

We thank CCMB COVID-19 testing facility, especially Karthik Bhardwaj and Archana Bhardwaj Siva for the access of viral VTMs. We thank Haripriya Parthasarathy for critical reading and her valuable suggestions. Special appreciation goes to Mohan Singh Moodu and Amit Kumar for their assistance with logistics.

## Supplementary Figures

**Supplementary Figure 1.**
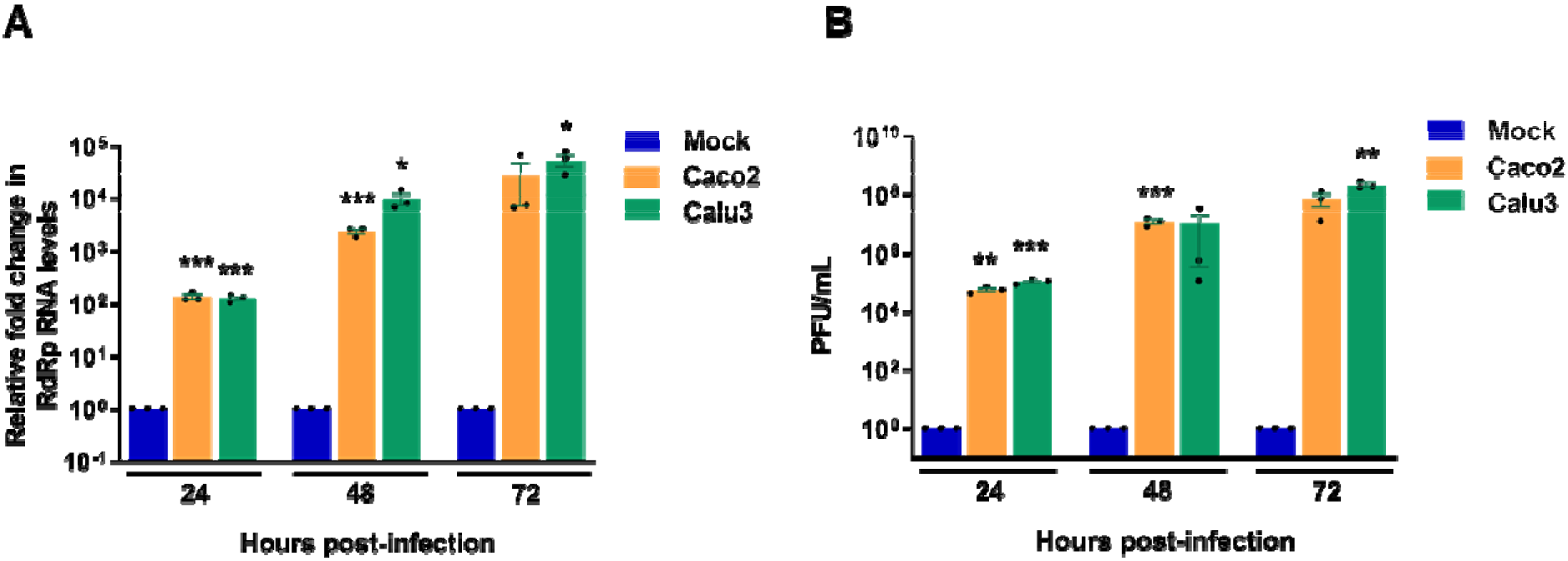
Permissivity analysis for SARS-CoV-2 in Calu3 and Caco2 cells. (A) Supernatants from Calu3 and Caco2 cultures infected with 1 MOI of B.1.1.8 variant isolate of SARS-CoV-2 for 24-, 48- or 72 h were analyzed by real-time qRT-PCR to measure the genomic RNA of SARS-CoV-2. The fold-changes over the mock-infected samples were calculated by ΔΔ-Ct method and plotted in the graph (B) Analysis of the infectious titers of SARS-CoV-2 in the supernatants generated in A, by PFA and represented in PFU/mL.

**Supplementary Figure 2.**
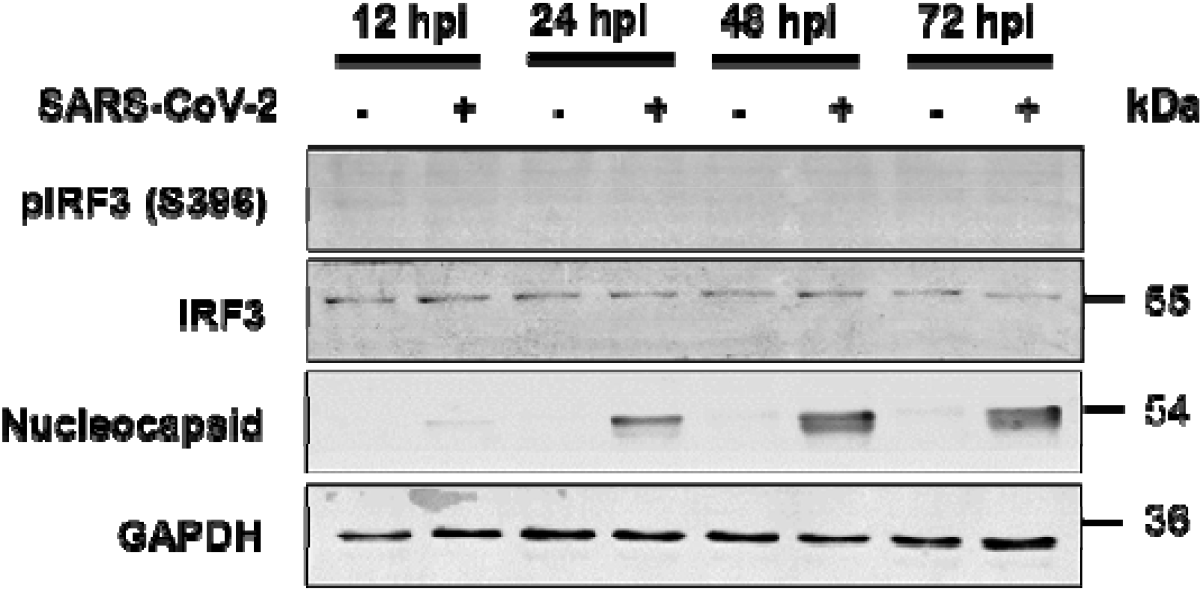
IRF3 phosphorylation is suppressed in SARS-CoV-2 infected Calu3 cells. Calu3 cells were infected with 1 MOI of B.1.1.8 variant isolate of SARS-CoV-2 for various time-points as indicated in the figures. Cells harvested were lyses and subjected to immunoblotting to detect viral N and phosphorylated IRF3.

**Supplementary Figure 3.**
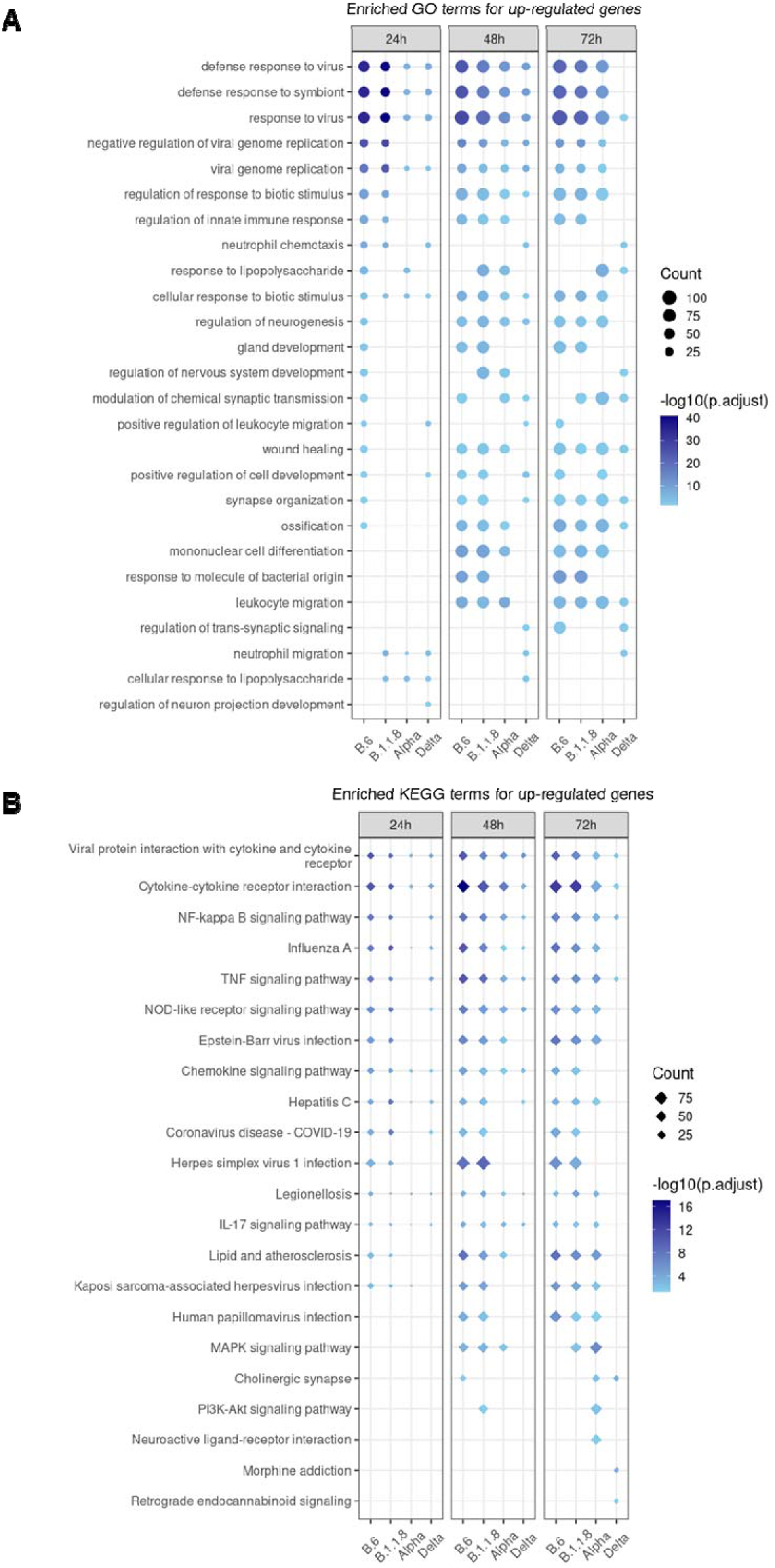
Enrichment analysis representing the Enriched GO (circles) and KEGG (diamond) terms for up-regulated DEGs for each variant-infected samples at each time-points. Size of the dot represents the number of DEGs in the enriched term and the intensity of the color represents the -log10 (adjusted p-value).

**Supplementary Figure 4.**
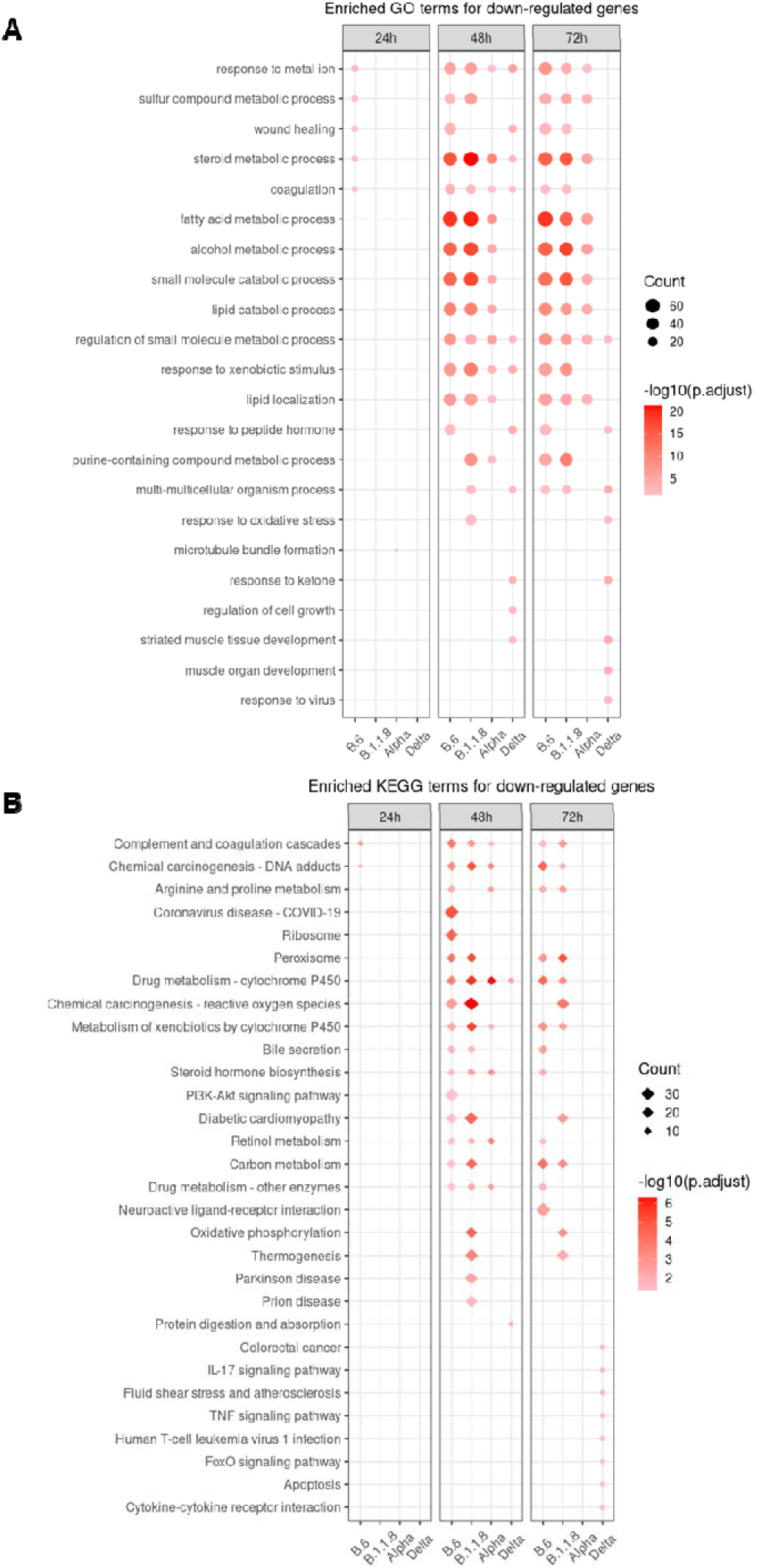
(A and B) Enrichment analysis representing the Enriched GO (circles) and KEGG (diamond) terms for down-regulated DEGs for each variant-infected samples at each time-points. Size of the dot represents the number of DEGs in the enriched term and the intensity of the color represents the -log10 (adjusted p-value).

**Supplementary Figure 5.**
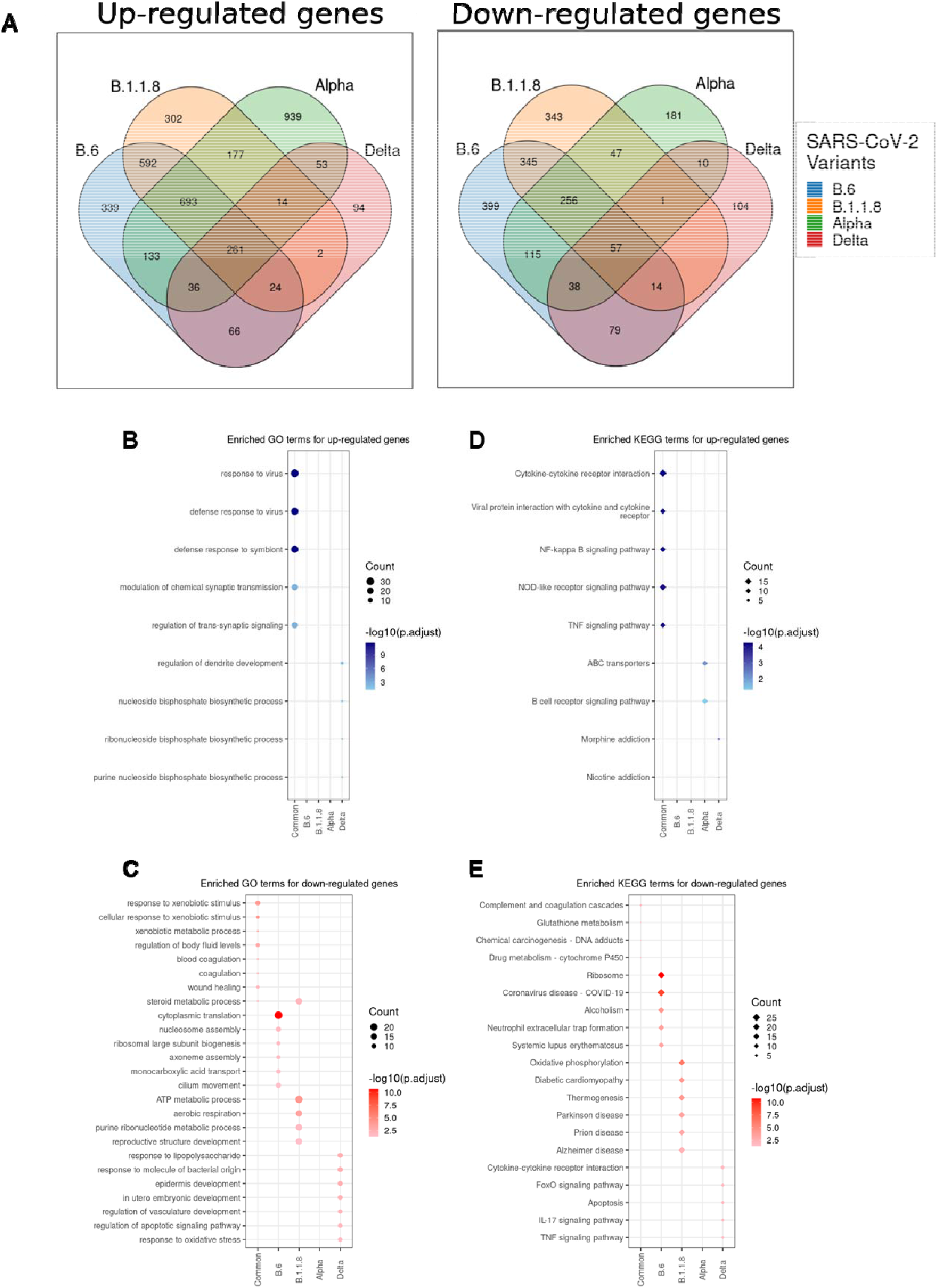
Analysis of the overlapping and unique DEGs from individually infected samples. (A) Venn diagram showing the common genes that were differentially regulated by all the four variants, as well as unique genes from each individual infections, for both up-regulated and down-regulated sets. DEGs were pooled from all time-points for each variant sample and used in the analysis. (B-E) GO and KEGG enrichment analysis of the common and unique up and down regulated DEGs from each infected samples

**Supplementary Figure 6.**
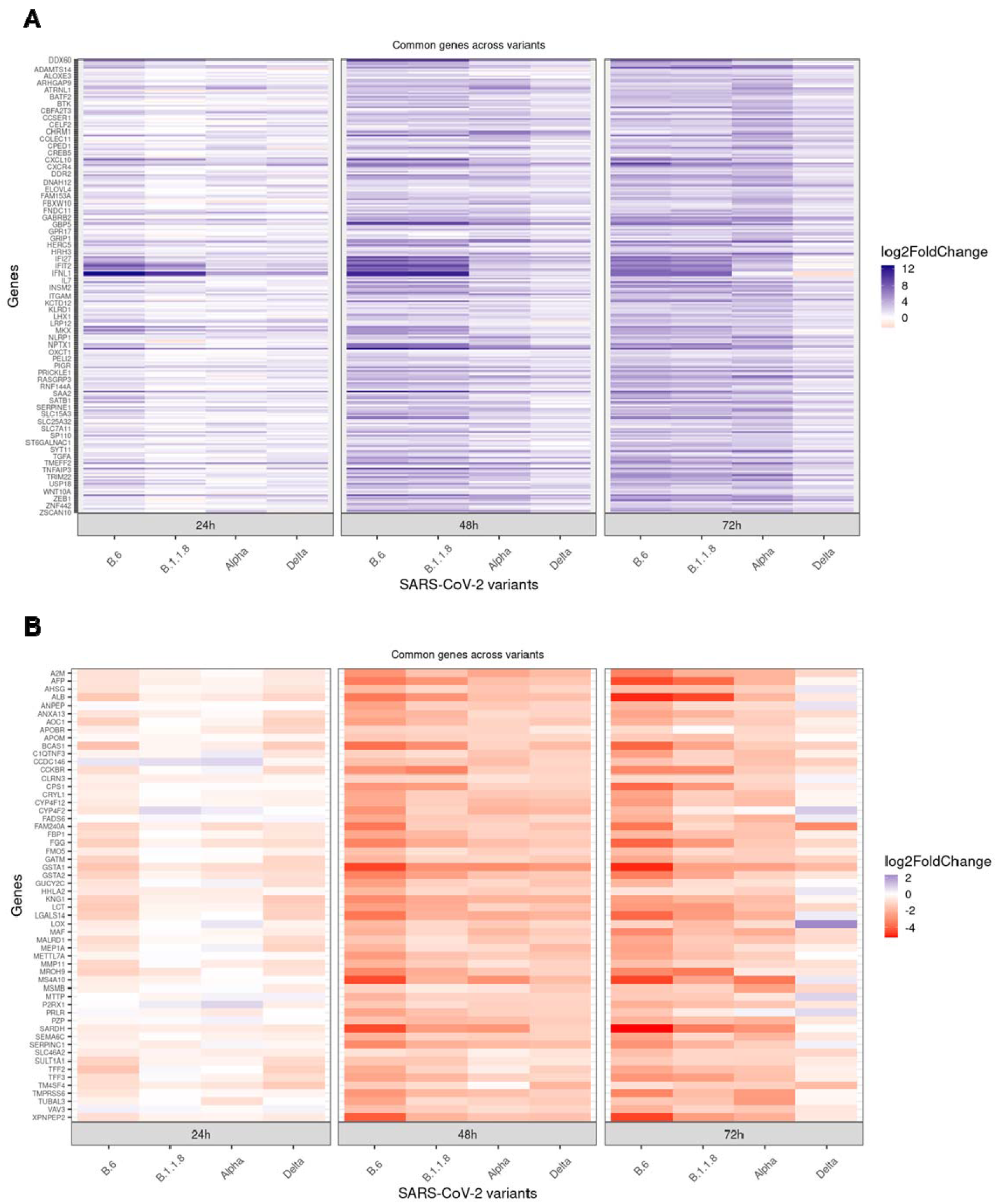
Heat-maps demonstrating the log2 fold change of (A) 261 up-regulated and (B) 57 down-regulated genes, common across the four variant infections as shown in Figure 5A. The lists of genes were generated from the common pool representing DEGs from all time-points as shown in Supplementary Figure 5A.

**Supplementary Figure 7.**
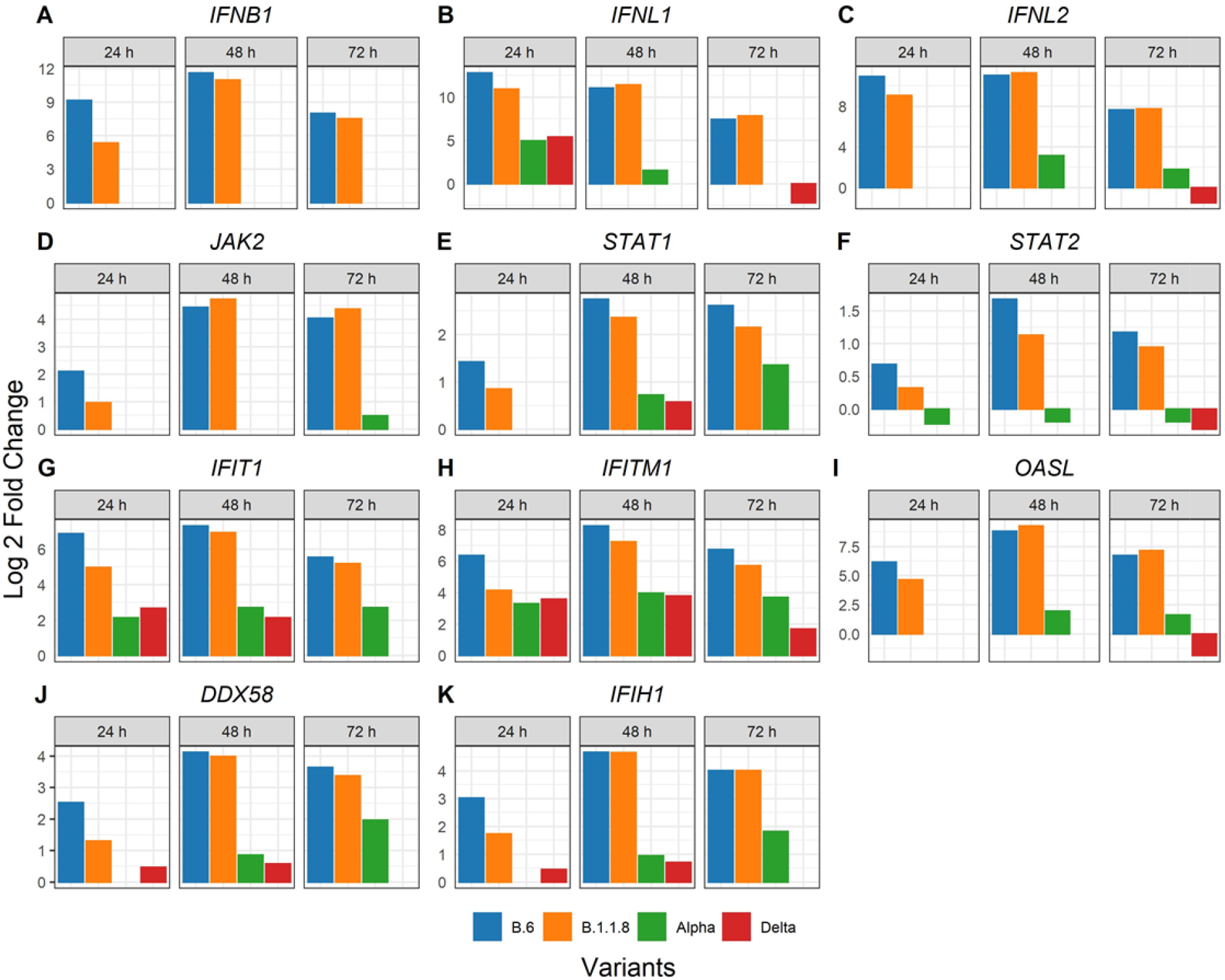
Bar-graphs demonstrating the differential expression of select genes of importance from type-I and type-III IFN pathways. The graphs were generated from the *p*-value adjusted list and are statistically significant.

**Supplementary Figure 8.**
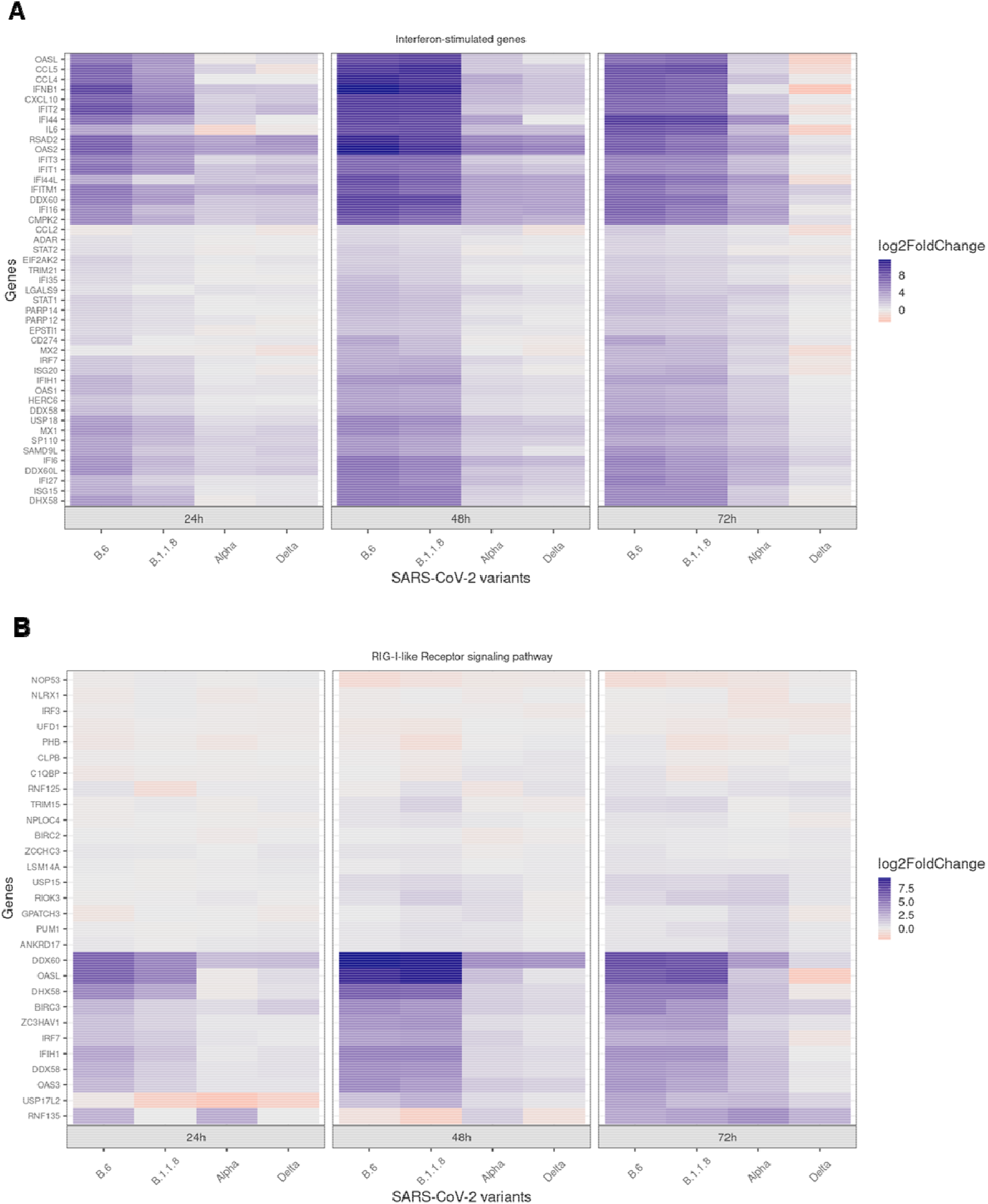
(A) Heat-map demonstrating the differential expression of ISGs in response to the variant infection at specified time-intervals. (B) Heat-map demonstrating the differential expression of genes classified under RLR pathway in response to the variant infection at specified time-intervals.

**Supplementary Figure 9.**
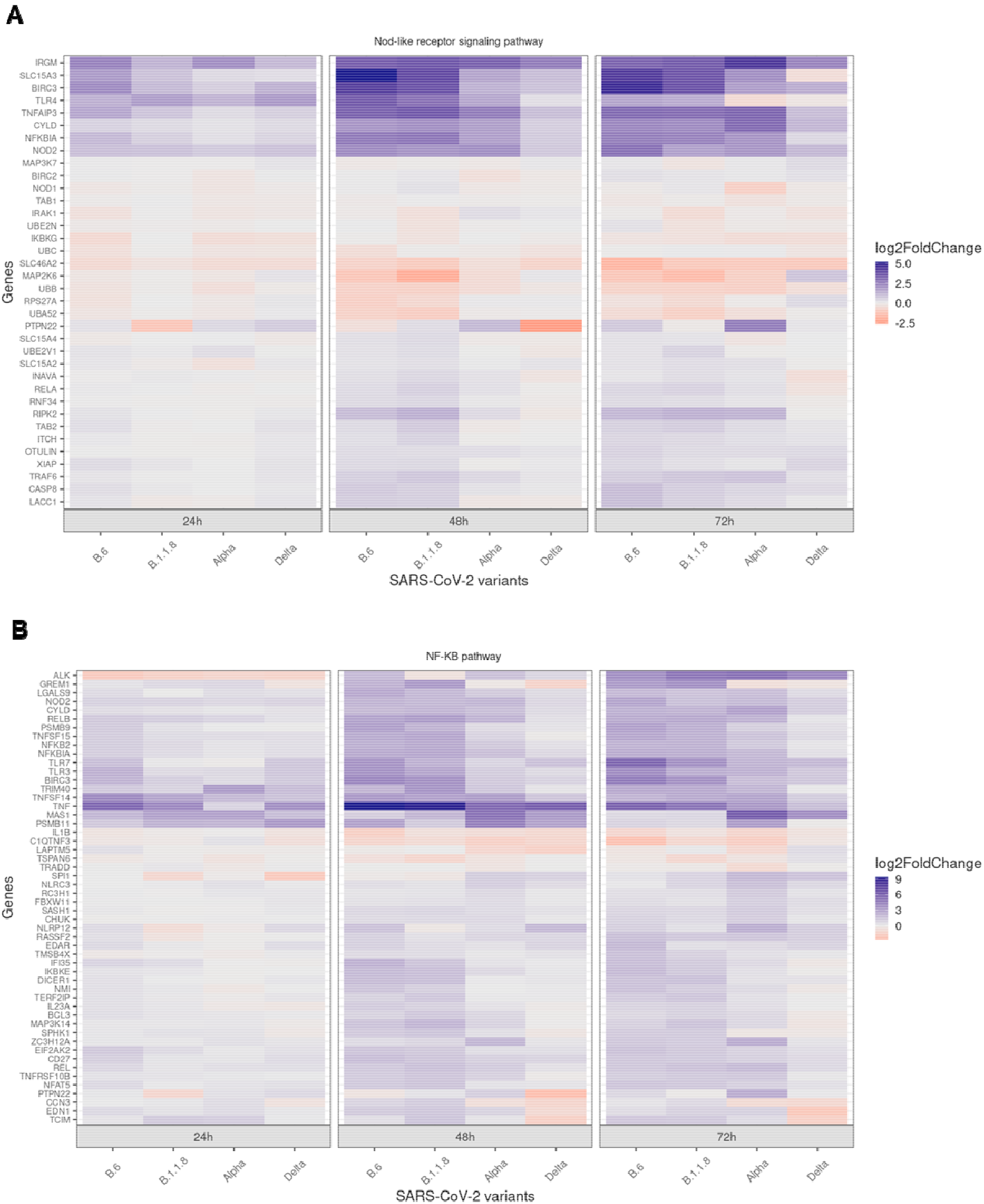
Heat-map demonstrating the differential expression of genes classified under NLR pathway in response to the variant infection at specified time-intervals. (B) Heat-map demonstrating the differential expression of genes classified under NF-κB pathway in response to the variant infection at specified time-intervals.

**Supplementary Figure 10.**
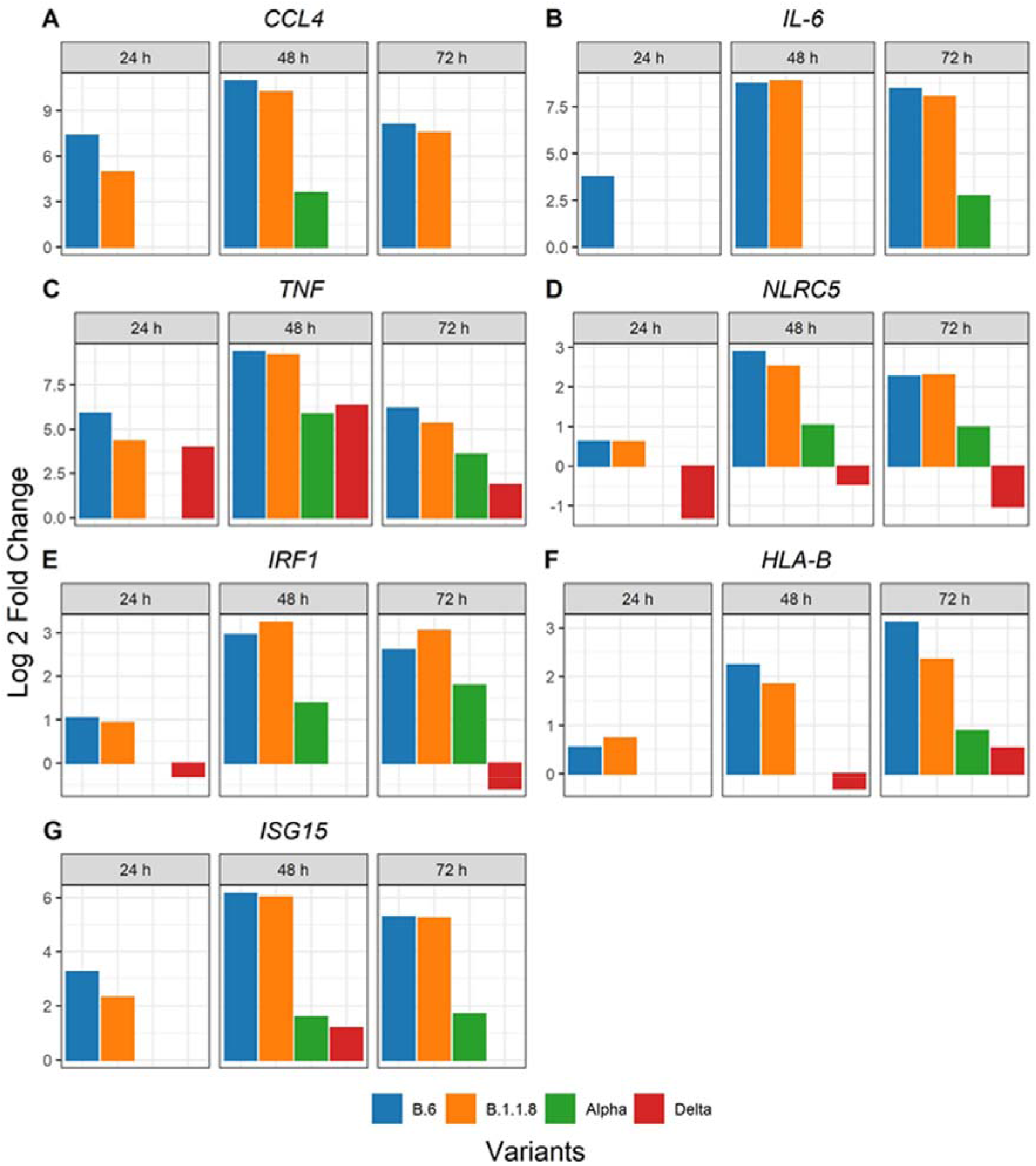
Bar-graphs demonstrating the differential expression of select genes (A-F) participating in antigen presentation, and regulation of interferon pathway (G, H). The graphs were generated from the *p*-value adjusted list and are statistically significant.

**Supplementary Figure 11.**
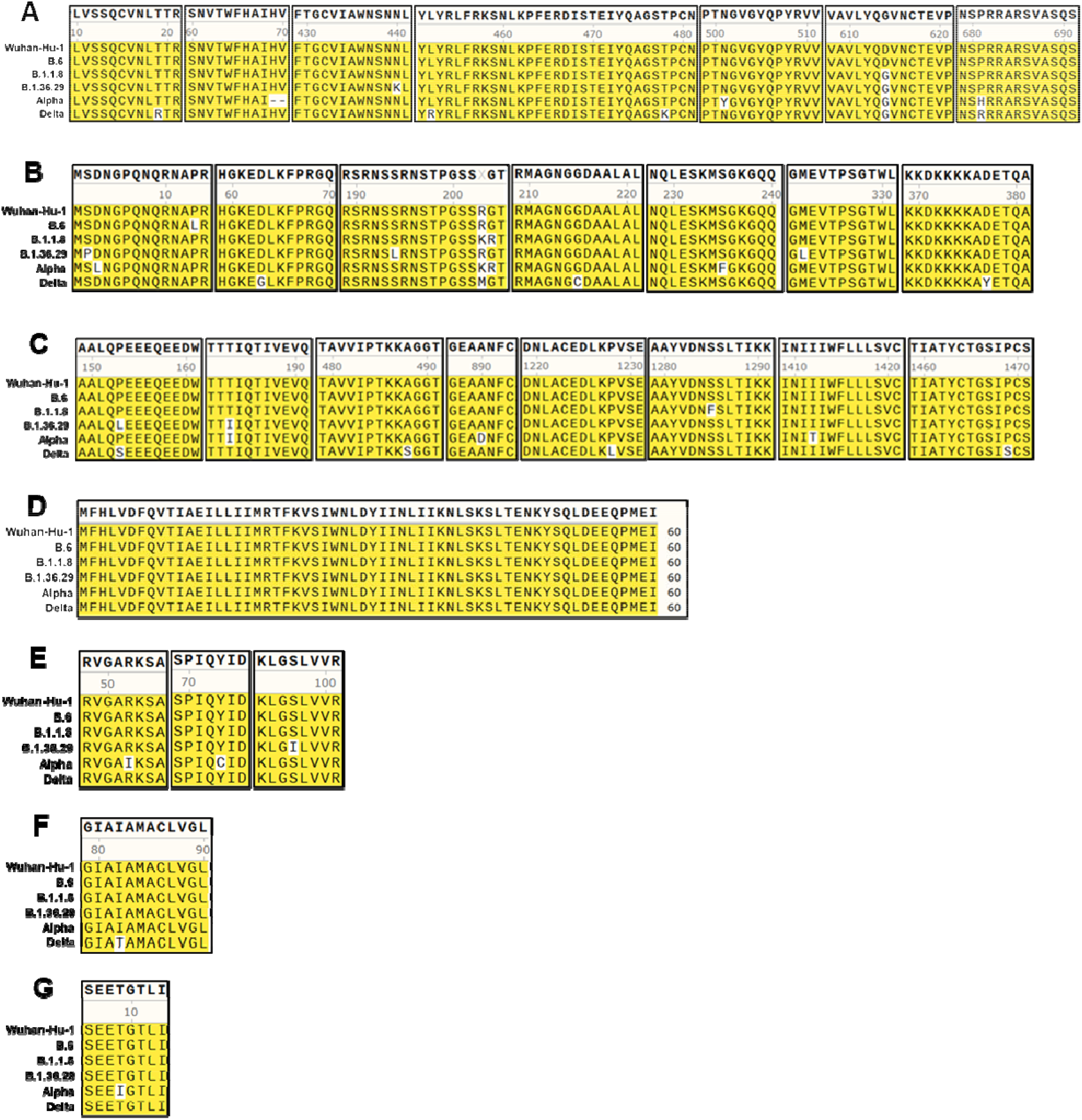
Alignment of regions of various SARS-CoV-2 polypeptide sequences from the variants used in this study. (A: Spike; B: Nucleocapsid; C: Nsp3; D: ORF6; E: ORF8; F: Membrane; and G: Envelope)

